# Aligning with the predecessors and counterarguments: A systematic review of the anatomical correlates for the newly discovered meningeal layer in the existing literature

**DOI:** 10.1101/2024.02.10.579742

**Authors:** Ashutosh Kumar, Rajesh Kumar, Ravi K. Narayan, Banshi Nath, Ashok K. Datusalia, Ashok K. Rastogi, Rakesh K. Jha, Pankaj Kumar, Vikas Pareek, Pranav Prasoon, Muneeb A. Faiq, Prabhat Agrawal, Surya Nandan Prasad, Chiman Kumari, Adil Asghar

## Abstract

A recent study reported the existence of a subarachnoid lymphatic-like membrane (SLYM) –an intermediate leptomeningeal layer between the arachnoid and pia mater in mice and human brains – dividing the subarachnoid space (SAS) into two functional compartments. Despite being a macroscopic structure, how it missed detection in previous studies is surprising. We systematically reviewed the published reports in animals and humans to explore whether prior descriptions of this meningeal layer have existed. An electronic search was conducted in PubMed/Medline, EMBASE, Google Scholar, Science Direct, and Web of Science databases using combinations of MeSH terms and keywords with Boolean operators from inception until 31^st^ Dec 2023. We found at least eight studies that provided structural evidence of an intermediate leptomeningeal layer in the brain or spinal cord. However, unequivocal descriptions for this layer all along the central nervous system were scarce. Obscured names were used to describe it, i.e., the epipial layer, intermediate meningeal layer, intermediate lamella, and outer pial layer. Its microscopic/ultrastructural details closely resembled the SLYM. Further, we examined the counterarguments in current literature that are skeptical of this layer’s existence. Considering the significant physiological/clinical implications, exploring further structural and functional details of the new meningeal layer is a need of the hour.

## 1. Introduction

**T**he meningeal coverings create a protective barrier around the brain and spinal cord. Their arrangement and architecture are crucial in the health and disease of the central nervous system (CNS). As per traditional knowledge, there are three layers in meninges, outer to inner: dura mater, arachnoid mater, and pia mater (Standring, 2021).

The dura mater outlines the cranial cavity and is an independent layer. However, the arachnoid and pia create a closed compartment along the brain and spinal cord— the subarachnoid space (SAS) filled with cerebrospinal fluid (CSF). The SAS is traversed by the numerous ‘arachnoid trabeculae’ attached to the arachnoid and pia mater. Recently, Møllgård *et al*. reported the existence of a new leptomeningeal layer in mice and human brains between arachnoid and pia, dividing the subarachnoid space into two functional compartments (Møllgård et al., 2023). They described a ‘subarachnoid lymphatic-like membrane (SLYM)’ as a one to two-cell thick mesothelial membrane not allowing the passage of moieties more than one μm in size and three kilodaltons in weight (Møllgård et al., 2023). Thus, it divides the SAS containing CSF into two functional compartments (Fig. 1).

**Figure 1.**
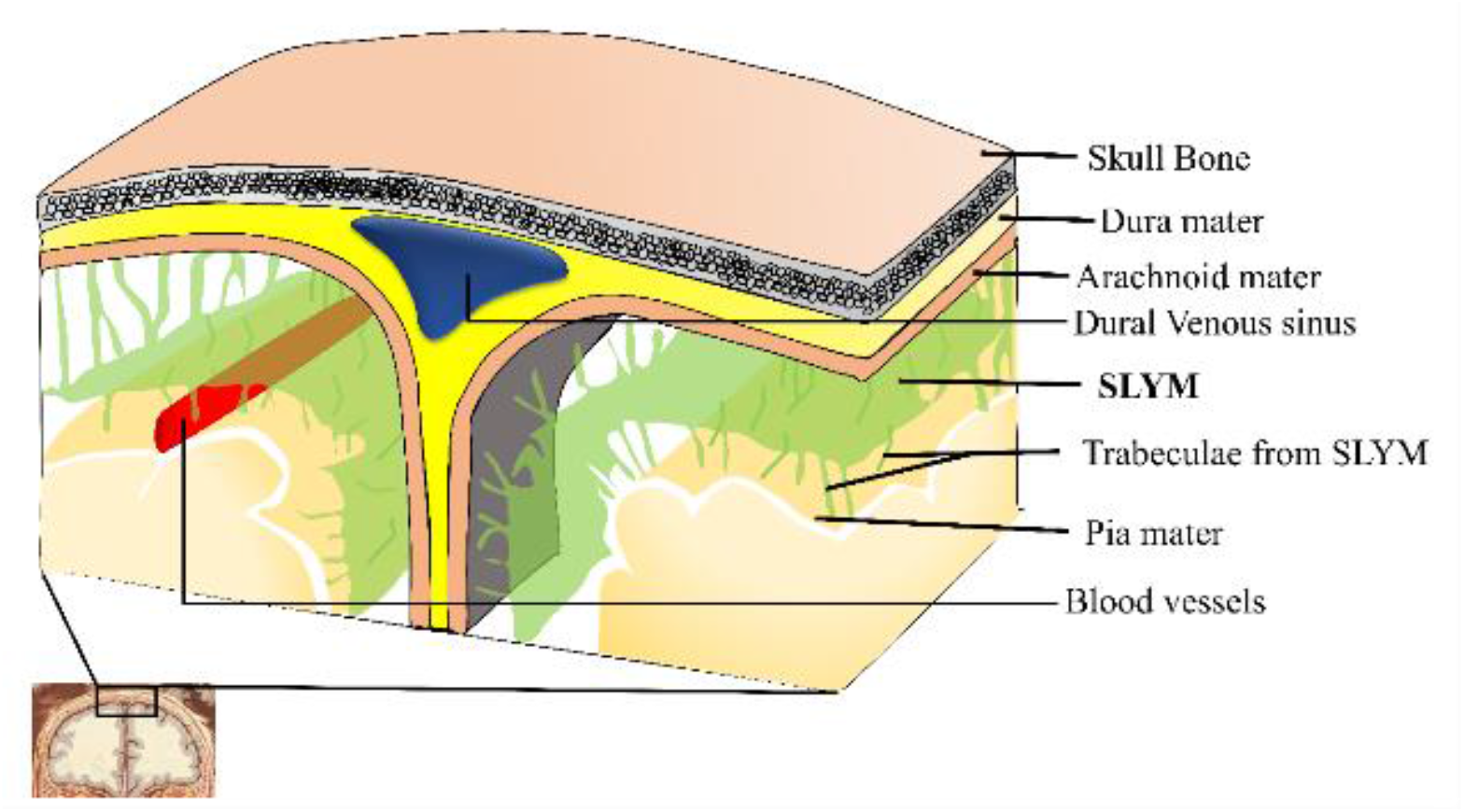
Revised meningeal arrangement in the human brain.

In this article, we looked for the prior indications of this new meningeal layer in the published literature and its alignment with the existing knowledge. Further, we examined the counterarguments in current literature that are skeptical of this layer’s existence.

## 2. Methods

### 2.1 Information sources and search strategy

We performed a scoping review of the published reports of meninges in CNS components of mammals, including humans, following the ‘Preferred Reporting Items for Systematic Reviews and Meta-Analyses (PRISMA)’ guidelines, 2020 (http://www.prisma-statement.org/). The online databases, including PubMed/Medline, EMBASE, Google Scholar, Science Direct, and Web of Science, were searched for the original studies using multiple combinations of controlled vocabulary terms and free keywords with Boolean operators. Advanced search builder was used for PubMed, applying all fields to widen the search. Further, a more specialized systematic search with targeted terms involving physiological and clinical terms was performed to discuss the new meningeal layer’s physiological and clinical significance/relevance.

### 2.2 Keywords and Search Strings

### MeSH terms

(meninges, dura mater, arachnoid, pia mater, cerebrospinal fluid, blood-brain barrier, anatomy, humans, primates, brain, spinal cord, central nervous system, subarachnoid space, magnetic resonance imaging, microscopy, electron, scanning, transmission, ultrasonography, computed tomography, lymphatic vessels, glymphatic system, growth and development, inflammation)

### Free keywords

subarachnoid lymphatic like membrane (SLYM), epipial layer, inner pial layer, outer pial layer, circulation, blood-CSF barrier, dissection, trabeculae, layers, clinical significance, immune cells, mesothelium, ligamentum denticulatum, linea splendens, filum terminale etc. For search strings see Supplementary file 1.

### 2.3 Eligibility criteria

Original reports of macroscopic, microscopic/ ultrastructural, neuroimaging, and clinical studies of brain or spinal meninges in animals (mammals only) and humans were included. The studies published up to 31^st^ Dec 2023 were eligible without language restrictions. The case reports, commentary, and newsletters were excluded.

### 2.4 Data selection, extraction, and synthesis

Full articles reporting original details on the meninges were included in the study. Data collection was performed in two steps. In the first step, the titles and abstracts were screened. The articles in languages other than English were translated using Google Translate. The studies did not meet the eligibility criteria, and the duplicates were excluded. In the second step, full-text articles were assessed for the included studies. Reasons for excluding any article at the second step were recorded as given in Fig. S1 (Supplementary file 2). No prioritization approach was adopted during the search to avoid a selection bias. Citation details and abstracts of all retrieved studies were downloaded into the Mendeley bibliography manager for the record. Three authors independently completed data collection. Two independent investigators performed the quality assessment of the included articles using the ‘Anatomical Quality Assessment (AQUA)’ tool for the anatomical studies included in meta-analyses and systematic reviews, as given in Table S1 (Supplementary file 3) (Henry et al., 2017).

All co-authors contributed to and reviewed the qualitative analysis. Mutual discussions resolved any investigative disagreement. The final inferences were made based on the qualitative synthesis from the collected data. No quantitative analysis or statistical testing was performed.

## 3 Results

A total of 1060 articles were identified. After excluding the irrelevant entries and duplicates, 50 research articles were selected for the title and abstract screening, out of which 42 were excluded for not fulfilling the inclusion criteria (PRISMA 2020 flow diagram, Fig. S1). Only eight articles were eligible for the analysis that directly mentioned and provided structural details of an intermediate leptomeningeal layer in spinal or brain SAS. The key observations from these studies are presented in Table 1.

**Table 1.** A chronological description of the intermediate leptomeningeal layer in the mammalian central nervous system’s subarachnoid space (SAS) in published literature.

| Serial N. | Author | Objectives | Key observations |
| --- | --- | --- | --- |
| 1. | Key and Retzius, 1875 | A descriptive study of pia mater in humans. | • *Epipial layer* in the spinal cord specimens from adult humans.<br>• A meshwork of collagenous fibers with a superficial irregular but deeper circular arrangement. |
| 2. | Millen and Woollam, 1961 | A descriptive study of pia mater in humans. | • *Epipial layer* in the cadaveric specimen of the spinal cord from adult and fetal humans<br>• Thick on the spinal cord's sides and anterior median fissure<br>• Formed ligamentum denticulatum, linea splendens, and filum terminale. |
| 3. | Krisch *et al.*, 1983 | A comparative analysis of the meningeal compartments of the median eminence and the cortex in rats. | • Four compartments were described around the cortex: 1. The intercellular compartment: comprised of the intercellular clefts of the neuropil and leptomeninges. 2. pial space. 3. arachnoid space. 4. intercellular clefts of the dura. |
| 4. | Krisch *et al.*, 1984 | To study compartments and perivascular arrangement of the meninges covering the cerebral cortex of the rat. | • *Intermediate lamella* in SAS of adult rat and rabbit brains<br>• This intermediate lamella comprised a single-cell thick outer pial layer enjoined with a similar thickness inner arachnoid layer.<br>• Both layers of the intermediate lamella continued as the perivascular sheath around the vessels. |
| 5. | Nicholas and Weller, 1988 | A light and scanning electron microscopy study of spinal meninges in humans. | • *Intermediate meningeal layer* in the cadaveric spinal cord specimen from adult humans.<br>• It comprised collagen fiber bundles arranged in lattices forming fenestrated sheets, thus giving free passage to the cerebrospinal fluid (CSF). |
| 6. | Angelov and Vasilev, 1989 | To study the morphogenesis of rat cranial meninges. | • *Outer pial layer* in developing rat brain. |
| 7. | Kurucz et al., 2013 a,b | Endoscopic anatomical study of the arachnoid architecture on the base of the skull | • Described inner arachnoid membranes around the brain parts in anterior, middle, and posterior cranial fossae in fresh human cadavers. |
| 8. | Mestre *et al.*, 2022 | To study perivascular spaces and associated degenerative changes with aging and Alzheimer's disease model in mice brain. | • *Epipial layer* in adult mice brain.<br>• Epipial layer cells identified as the reticular fibroblasts displaying three generations of cytoplasmic processes.<br>• The cytoplasmic processes adjoined with the help of junctional complexes (i.e., tight junctions), forming a microporous meshwork. |
|  |  |  | • Formed perivascular sheaths around the cerebral arteries that facilitated CSF tracer accumulation, indicating its role in CSF circulation.<br>• Degenerative changes in the epipial vascular sheath in mice models of aging and Alzheimer's disease. |

## 4 Discussion

### 4.1 Macroscopic and microscopic/ultrastructural anatomy

The recent study by Møllgård *et al*. uniquely claimed to discover a new leptomeningeal layer in mice and human brains using immunohistology, *in vivo* two-photon microscopy, and CSF tracer-based methods (Møllgård et al., 2023). The existing literature principally details the architecture of only three meningeal layers, i.e., dura, arachnoid, and pia mater. However, occasional studies in animals and humans provide indications of an intermediate leptomeningeal layer in the SAS of the brain and spinal cord (Table 1)(Angelov & Vasilev, 1989; Key, A., and Retzius, 1875; Krisch et al., 1984; Mestre et al., 2022; Millen & Woollam, 1961; Nicholas & Weller, 1988).

The macroscopic description of meninges in history can be traced back to as early as 1600 BC, in the Edwin Smith Papyrus from ancient Egypt (Adeeb et al., 2013). The given macroscopic descriptions of meninges in Papyrus perhaps indicated the dura mater (Adeeb et al., 2013). In the fourth century BC, the Greek philosopher Aristotle, based on animal dissections, described two layers of meninges: the stronger one near the skull and the delicate one covering the brain, indicative of dura mater and pia mater, respectively (Adeeb et al., 2013). Although the presence of arachnoid mater between dura and pia mater was first described as early as the third century BC by Herophoilus, its detailed descriptions were given much later by Frederick Ruysch, a Dutch anatomist, in 1699 (Adeeb et al.,2013). Thus, a conventional order of referring outer to inner meningeal layers was set in practice as dura, arachnoid, and pia mater, and modern studies provided extensive structural details for each of these layers (Coles et al., 2017).

The discovery of the SLYM between the arachnoid and pia changes the set order of the meningeal layers. The outermost meningeal layer—dura mater *(dura-* tough, *mater-mother)* is a thick, dense collagenous membrane called pachymeninx (*patchy*-thick). In contrast, the intermediate layer, the arachnoid mater, and the innermost layer, the pia mater, are called leptomeninges *(lepto-* thin) (Standring, 2021). Now, the SLYM is a new entrant in the category of leptomeninges. Being positioned between the arachnoid and pia mater in SAS, it has to be considered the intermediate leptomeningeal layer. In the revised hierarchy of the meninges, it ought to be recognized as the third meningeal layer from the outside, shifting the pia mater to the fourth.

The published literature’s foremost mention of an intermediate leptomeningeal layer in human SAS limited its presence to the spinal cord. As early as 1875, Key and Retzius described the inner leptomeninx as pia intima and epipial tissue. Pia intima directly posed on the nervous tissue and resembled the current description of pia mater (Key, A., and Retzius, 1875). The epipial tissue layer was described as a meshwork of collagenous fibers with a superficial, irregular, but deeper circular arrangement. Key and Retzius’s descriptions (Key, A., and Retzius, 1875) did not receive general support until 1961 when Millen and Woollam (Millen & Woollam, 1961) reconfirmed them in normal human adult and fetal spinal cords. Millen and Woollam noted that the epipial tissue layer was thick on the sides of the spinal cord and the anterior median fissure. On the sides, it formed the collagenous core of the ligamentum denticulatum. In the anterior median fissure, thickened epipial tissue formed linea splendens that enclosed anterior spinal vessels. The epipial tissue also dipped into the depth of the anterior median fissure (Millen & Woollam, 1961).

Moreover, the filum terminale, the tubular extension from the terminal part of the spinal cord, was also described as formed of the epipial tissue layer. The epipial tissue layer was noted as a continuous sheet and only interrupted at the sites of entry and exit of the spinal nerve rootlets, forming a round margin around the nerves. The subarachnoid vessels lay between the epipial tissue layer strands before entering the spinal cord’s substance (Millen & Woollam, 1961). The authors could not find its presence in the brain beyond the medulla oblongata. Later, in 1988, Nicholas and Weller described the presence of an intermediate meningeal layer between the arachnoid and pia in the human spinal cord using scanning electron microscopy (SEM) (Nicholas & Weller, 1988). Their portrayal of the intermediate meningeal layer in the spinal cord resembled the epipial tissue layer described by Key and Retzius (Key, A., and Retzius, 1875) and Millen and Woollam (Millen & Woollam, 1961). Nicholas and Weller also described the intermediate meningeal layer contributing to the ligamentum denticulatum and enveloping anterior spinal vessels at the anterior median fissure.

Interestingly, the epipial tissue layer (Millen & Woollam, 1961) or the intermediate meningeal layer (Nicholas & Weller, 1988) was described by the authors as the collagen fiber bundles arranged in lattices forming fenestrated sheets, thus giving free passage to the CSF. Also, there were no mentions of the leptomeningeal cells lining them. The permeability for CSF and absence of leptomeningeal cell lining in these descriptions contrast the SLYM’s portrayals presented by Møllgård *et al*. However, Møllgård *et al*.’s descriptions were limited to the brain as they did not give any data from the spinal cord.

Unfortunately, the historical reports of the epipial tissue layer (Millen & Woollam, 1961) or intermediate meningeal layer (Nicholas & Weller, 1988) in the spinal cord were little followed in the later studies. Millen and Woollam’s descriptions of the epipial tissue layer in the formation of ligamentum denticulatum, linea splendens, and filum terminale (Millen & Woollam, 1961) were also obscured as these structures have been conventionally described as the modifications of the pia mater. We discussed Nicholas and Weller’s SEM findings in detail under the sub-section ‘Hidden indications from SEM data.’

The foremost indication of the intermediate meningeal layer in brain SAS is observable in a couple of studies in 1983-84 in rats and rabbits by Krisch *et al* (Krisch et al., 1983, 1984). The authors used IHC and EM to show the presence of a two-cell thick intermediate lamella that compartmentalized the brain SAS into pial and arachnoid spaces (Fig. 2).

**Figure 2.**
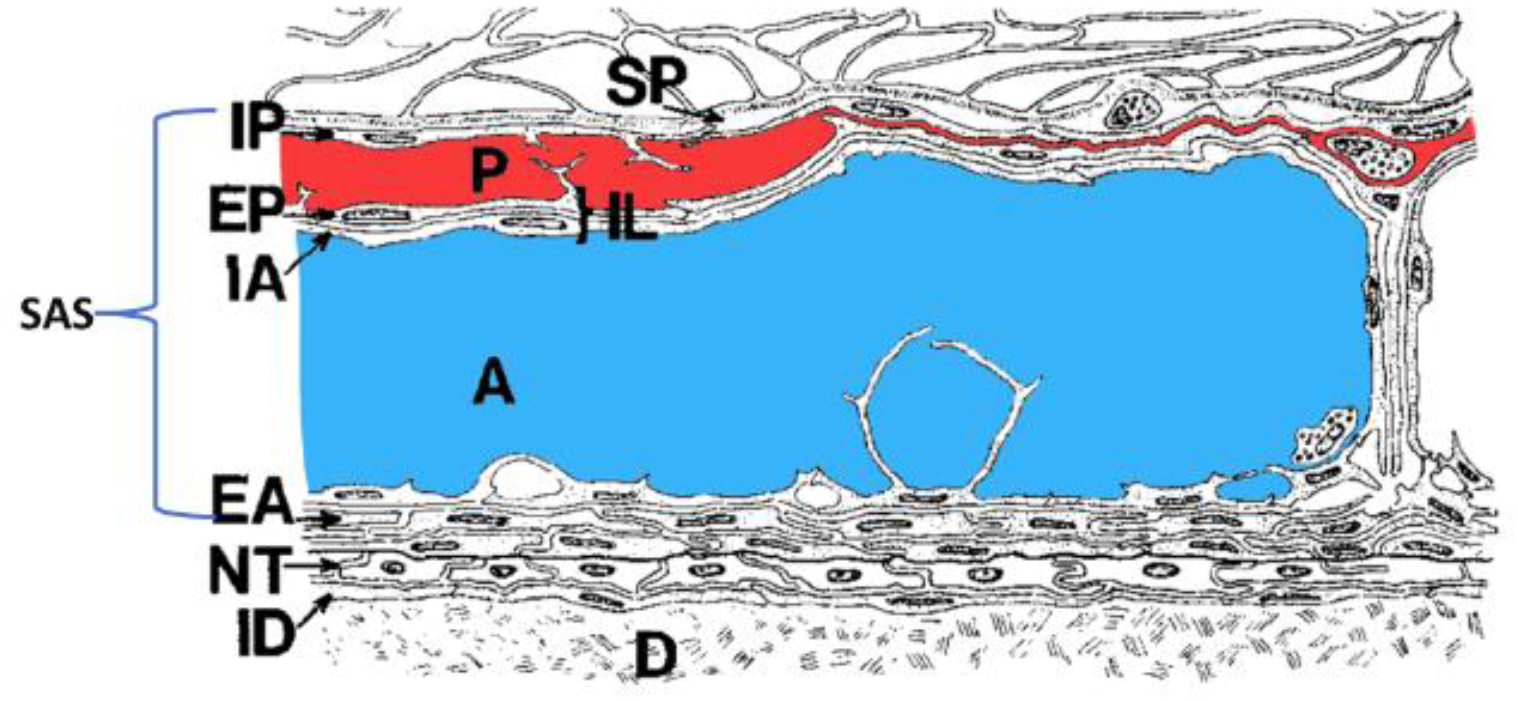
Schematic representation of the meningeal arrangement around rat cortex originally demonstrated by Krisch et al., 1983 in TEM imaging. [A slightly modified schematic representation of meningeal arrangement around rat cortex originally depicted by Krisch *et al*., 1983 based on his TEM imaging observations. A two-cell thick intermediate lamella (IL) compartmentalizes the brain SAS into pial (P, labeled in red) and arachnoid (A, labeled in blue) spaces. The IL comprises of a single-cell thick outer pial layer (EP) enjoined with a similar thickness inner arachnoid layer (IA). The pial space is extremely narrow at some sites and is again reopened. Abbreviations: SP: subpial space; IP: inner pial layer; P: pial space; EP: outer pial layer; IA: inner arachnoid layer. IL: intermediate lamella; A: arachnoid space; EA: outer arachnoid layer; NT: neurothelium; ID: inner dural layer; D: dura mater. **Reproduced with permission from Krisch *et al***., **1983**.]

This intermediate lamella comprised a single-cell thick outer pial layer enjoined with a similar thickness inner arachnoid layer. Both layers of the intermediate lamella were also continued as the perivascular sheath around the vessels. The pial space became extremely narrow at some sites and again reopened at other sites. Replacing the conventional description of two leptomeningeal layers, the authors proposed “inner, intermediate”, and “outer leptomeninges,” encompassing SAS’s inner and outer leptomeningeal spaces.

When authors injected horse radish peroxidase (HRP) in either the pial or arachnoid space, the dye was limited to that space only, indicating the impermeability of the intermediate lamella bordering these spaces (Krisch et al., 1984). Interestingly, intracerebral injection of HRP labeled the intercellular clefts of the intermediate lamella as well as the outer arachnoid layer but not the pial or arachnoid spaces, indicating unexplained channels of communications between intercellular clefts across the leptomeningeal layers and brain parenchyma (Krisch et al., 1984).

In recent years, Kurucz *et al*., in consecutive neuro endoscopic studies in fresh human cadavers, described inner arachnoid membranes in brain SAS that match the positional description given for SLYM. (Kurucz et al., 2013a, 2013b). The authors demonstrated that the inner arachnoid membrane was distinct from the outer arachnoid, with delicate trabeculae separating the layers. However, as a significant limitation, they provided only localized and fragmented descriptions, and it was unclear if the inner arachnoid membrane presented as a single and continuous layer along the neural axis. The SAS was more complicated at specific locations, which lodged large vessels and anastomotic channels, such as in the Sylvian fissure, interpeduncular, carotid, and pontine cisterns. Multiple inner arachnoid layers were shown at these places, some sieved and appearing as trabecular networks (Kurucz et al., 2013a, 2013b)..

Interestingly, immediately before Møllgård *et al*. discovered SLYM, another study *by* Mestre *et al*. described the epipial tissue layer in mice brains (Mestre et al., 2022). The authors presented an epipial layer of the pia mater in the cerebral cortex using advanced light and electron microscopy combined with routine and immunohistological stainings and CSF tracers methods. Mestre *et al*.’s descriptions closely resembled SLYM. However, their study primarily focused on delineating the contribution of the epipial layer in forming a perivascular sheath and lacked detailed exploration along the brain surfaces. Keeping with the earlier descriptions of the epipial tissue layer in the spinal cord, the authors seemed to consider it a component of the pia mater, as they did not examine its immunophenotypic differences from the arachnoid and pia mater (Mestre et al., 2022). The pial and epipial layer cells were identified as the reticular fibroblasts, displaying three generations of cytoplasmic processes. The cytoplasmic processes of the reticular cells were shown adjoined with the help of junctional complexes (i.e., tight junctions), forming a microporous meshwork. They further showed that the epipial layer formed perivascular sheaths around the cerebral arteries that facilitated CSF tracer accumulation, indicating its role in CSF circulation. The authors also showed degenerative changes in the epipial vascular sheath in mice models of aging and Alzheimer’s disease (Mestre et al., 2022). As an exception, Mestre *et al*.*’s* description of the epipial layer also showed a deviation from that of SLYM by Møllgård *et al*. as the later demonstrated absence of tight junctions (Møllgård et al., 2023). However, Møllgård *et al*.*’s* observations were based on the expression of a tight junction marker, claudin-11, that differentiated SLYM from arachnoid mater, which expressed it abundantly (Møllgård et al., 2023). To resolve these differences, an EM-based exploration of SLYM’s junctional complexes is advisable.

Macroscopically, all three earlier known meningeal layers completely cover the CNS. While the outer two layers are closely apposed, the inner two enclose a fluid-filled trabeculated space— SAS (Coles et al., 2017; Susan Standring, 2021). The dura mater outlines the cranial cavity and vertebral canal. The arachnoid mater maintains a loose apposition to the brain surface; however, it bridges over the sulci, cisterns, and between the parts of the brain. In contrast, the pia mater is tightly apposed to the brain surface and follows it into the sulci and cisterns (Coles et al., 2017; Susan Standring, 2021). The SAS is just a capillary layer thick at the gyri and flattened surface in CNS, but it is deep and contains a substantial amount of CSF in the sulci and cisterns (Coles et al., 2017; Susan Standring, 2021). Histologically, the arachnoid mater is thin and translucent—made of a few layers of flattened cells (Pawlina & Ross, 2020.). From the arachnoid mater arise the arachnoid trabeculae made of spongy connective tissue consisting of collagen fibers and fibroblasts (Coles et al., 2017; Pawlina & Ross, 2020.). Compared to the two outer layers, which include a substantial amount of fibers, the pia mater is a delicate, single-cell-layer membrane (Coles et al., 2017; Pawlina & Ross, 2020.). The basement membrane of the pia mater is fused with that of the glia limitans—the astrocytic sheath covering the brain parenchyma, with a nominal subpial space, thus closely adhering to the brain surface (Adeeb et al., 2013; Dasgupta & Jeong, 2019; Susan Standring, 2021).

Although no macroscopic examinations were conducted, patterns revealed from the histological images in Møllgård *et al*.’s study suggest that the SLYM is very thin (14.2 ± 0.5 μm in mice brains) but has a macroscopically visible structure (Møllgård et al., 2023). Like the arachnoid mater, it passes over the sulci. Thus, the inner compartment would be more profound at the sulci than the outer CSF compartment. However, its arrangement over the cisterns and fissures is still not documented. Although the SLYM seems present in the entire length of SAS, Møllgård *et al*. did not provide any data from the spinal cord. The vessels have been described to be present in the inner compartment. The authors showed in the histological sections that SLYM enwraps the vessels. However, whether it gets carried forward as a tubular sheath is unclear from their study (Møllgård et al., 2023). The studies that have explored its gross details and neurosurgical/dissection methods to show it macroscopically are in wait. Microscopically, it is only a one to two-cell thick membrane intermixed with loosely organized collagen fibers. Møllgård *et al*. described it as an impermeable membrane that does not allow passage of moieties larger than one μm and three kilodaltons, which limits the exchange of most of the peptides and proteins, such as amyloid-beta and tau. Thus, the new layer divides SAS into two functional compartments (Møllgård et al., 2023). Studies that can provide its ultrastructural details are still lacking. Based on the histology and immunophenotypic characterization by Møllgård *et al*., SLYM closely resembles the lymphatic vessels (PDPN^+^ Prox^+^ Lyve^-^). It is immunophenotypically distinct from the arachnoid mater, pia mater, and arachnoid trabeculae. The leptomeningeal cells in SLYM lack tight or adherence junctions as they showed negative expressions of claudin-11 and E-cadherin (Møllgård et al., 2023).

The trabeculae were shown to anchor SLYM to the arachnoid mater above and the pia mater below. Møllgård *et al*. did not mention if there were distinct immunophenotypic differences between the trabeculae in the outer and inner compartments; hence, this remains to be explored (Møllgård et al., 2023). The subarachnoid blood vessels were shown primarily in the inner compartment between SLYM and the pia mater (Fig. 1) (Møllgård et al., 2023).

Unfortunately, no EM-based descriptions are currently available that can help differentiate SLYM ultrastructurally from other meningeal layers. Based on the histological features and permeability details presented by Møllgård et al., such as negative expressions of tight and adherence junction markers, claudin-11and E-cadherin, respectively, it can be speculated that its architecture is closer to the pia mater than arachnoid mater. In the EM, the arachnoid appears as a thick translucent sheet (about 200 μm in humans) composed of closely packed leptomeningeal cells joined by adherence junctions (i.e., desmosomes) and devoid of basement membranes (Saboori, 2021; Saboori & Sadegh, 2015; Roy O. Weller et al., 2018). Its outer barrier layer abuts the dura mater, where tight junctions join the cells. The inner layer of the arachnoid facing the CSF –the reticular layer— has a thin underlying layer of collagen that extends into trabeculae. A thin layer of leptomeningeal cells underlines that reticular layer and extends over the subarachnoid trabeculae. In comparison, the pia mater comprises a thin layer of leptomeningeal cells joined mainly by the gap junctions with the occasional presence of desmosomes (R. O. Weller, 2005; Roy O. Weller et al., 2018).

In addition to these structural details, the authors presented robust evidence in mice brains that SLYM may have a role in CNS immune responses as it harbored immune cells (CD45+ macrophages) that increased with aging and in the presence of inflammation (Møllgård et al., 2023).

### 4.2 Embryogenesis

The developmental evidence for an intermediate leptomeningeal is scarce in the literature. In a rare study in 1989, Angelov and Vasilev, using light and electron microscopy, described the presence of inner and outer pial layers in the embryonic rat brains enclosing blood vessels in between (Angelov & Vasilev, 1989). This description resembles the inner compartment of the SAS in the study by Møllgård *et al*., with inner and outer pial layers representing the pia mater and SLYM, respectively.

#### 4.2.1 Contradictions with proposed mesothelial derivation

In mammals, including humans, the primordial source for meninges development is the mesenchymal tissue surrounding the neural tube: the primary meninx (also called primitive meninx or meninx primitiva). The primary meninx is divided into a dense outer layer and an inner reticular layer. The inner layer is considered to be the meningeal mesenchyme. The meningeal primordium is further separated into pachymeninx (dura mater) and leptomeninx (arachnoid and pia mater) by the dural limiting layer. While pachymeninx contains longitudinally arranged fibroblasts, the leptomeninx is a meshwork of loosely organized cells. With further development, the leptomeninx undergoes cavitation to make the arachnoid trabeculae and the subarachnoid space bounded by the arachnoid and pia mater at its outer and inner limits, respectively (Batarfi et al., 2017; Dasgupta & Jeong, 2019; Schoenwolf GC, Bleyl SB, Brauer PR, 2021).

This primordial source also gives rise to the skull and the scalp. Further, the primary meninx also gives rise to a perineural vascular plexus, which will further develop into meningeal blood vessels. The histological observations suggested that the mesenchymal cells in the primitive meninx originate primarily from the neural crest cells (Batarfi et al., 2017; Dasgupta & Jeong, 2019; Schoenwolf GC, Bleyl SB, Brauer PR, 2021). The neural crest cells are, in turn, derived from the ectoderm at the margin of the neural tube following mesenchymal transformation (Schoenwolf GC, Bleyl SB, Brauer PR, 2021). Certain studies also suggested a contribution from the paraxial mesoderm in primitive meninx’s derivation (Couly et al., 1992; O’Rahilly & Müller, 1986). Notably, the paraxial mesoderm has an established contribution to the axial skeleton, including the head region (Tani et al., 2020); hence it is a likely contender for developing leptomeninges. However, evidence is scarce for the role of lateral plate or intermediate mesoderm in developing the axial skeleton and any meningeal layers.

Møllgård *et al*. described the SLYM as a mesothelium based on the immunohistochemical expression of podoplanin (PDPN), which is commonly expressed in the mesothelial membranes of the body cavities around the heart, lungs, and abdominal organs (Møllgård et al., 2023). Based on the PDPN expression, the authors compared the embryonic derivation of SLYM to the mesothelial lining of these body cavities, hinting at their shared embryonic development. These body cavities are embryologically derived from the intraembryonic coelom in the lateral plate mesoderm. If as such accepted, the proposed mesothelial derivation of the SLYM cognates it as a derivative of lateral plate mesoderm rather than paraxial mesoderm or neural crest cells. Thus, it contradicts the existing knowledge about the embryogenesis of the meningeal layers (Prummel et al., 2020; Schoenwolf GC, Bleyl SB, Brauer PR, 2021). Arguably, any such proposition will require a radical revision of human embryology and the evolution of chordates as it necessitates the presence of intraembryonic coelom (thus lateral plate mesoderm) inside the skull or vertebral column (Schoenwolf GC, Bleyl SB, Brauer PR, 2021).

Moreover, in stark contrast to the proposed single-layer structure in SLYM, mesothelial derivatives are double-layered, enclosing fluid-filled cavities (Schoenwolf GC, Bleyl SB, Brauer PR, 2021). Moreover, PDPN, used as a mesothelial marker, is not specific to intra-embryonic coelomic mesothelium. It is also expressed in derivatives of paraxial mesoderm, such as lymphatic vessels (Stone & Stainier, 2019), and intermediate mesoderm, such as podocytes in the glomerulus of the kidney (Tanaka et al., 2022). Moreover, it is also expressed in many non-mesodermal derivatives, such as type 1 pneumocytes and thymic medullary epithelial cells, which are endoderm derivatives (Astarita et al., 2012). Thus, PDPN alone only provides a critical lead but does not inform the mesodermal component from which SLYM is derived. The immunohistochemical examination of the markers for all mesodermal parts will be indispensable in establishing its embryonic derivation (Dasgupta & Jeong, 2019).

### 4.3 Counterarguments against the claims of a discovery

Odd with Møllgård *et al*.’s claim of breakthrough discovery, two recent studies presented strong reservations about the very existence of the SLYM (Mapunda et al., 2023; Pietilä et al., 2023). Pietilä *et al*., in a single-cell RNA-Sequencing (Sc-RNA seq) study of mouse meningeal fibroblasts, identified six different expression profiles (type 1-6) of leptomeningeal cells. Transgenic reporter labeling showed type 3 fibroblasts matched the Prox1+ cells in the arachnoid mater juxtaposed to the barrier cells (type 4 fibroblasts). They noted that Prox1+ cells are discontinuous, possibly intermixing with type 2 fibroblasts in the inner arachnoid layer. The inner arachnoid was identified distal to the barrier cell layer based on the specific ultrastructural features revealed in TEM. They further showed that the inner arachnoid connects with the barrier cell layer with the adherens junctions. Although the authors didn’t perform any immunostaining for Prox1+ cells in TEM, it was performed for Dpp4— a marker of the barrier cell layer. A localization study with Prox1 and Dpp4 in sequential sections may be needed to give more precise information on the proximity of Prox1+ cells with the barrier cell layer (Pietilä et al., 2023).

Moreover, Møllgård *et al*. didn’t describe the characterization of the fibroblasts lining the arachnoid trabeculae. Of note, the fibroblastic layers transversely traverse through the SAS, creating distinct compartments that can be conferred from the schematic representations of TEM images of mammalian brains in classic historical literature (Figs. 2&3) (Krisch et al., 1983, 1984; Nabeshima et al., 1975; Orlin et al., 1991). Although Møllgård *et al*. cite these precedences to corroborate their immunohistochemical findings, they performed no TEM imaging. A more detailed TEM imaging of the SAS with immunolabeling of the structures in context may be necessary to conclude the existence of the SLYM.

Another study by Mapunda *et al*. performed two-photon live imaging of injected fluorescent tracers in VE-cadherin-GFP reporter mice (Mapunda et al., 2023). The authors showed that VE-cadherin—a marker of the adherens junctions in the vascular endothelium— was abundantly expressed by leptomeningeal cells of the arachnoid and pia mater. They further observed that tracers injected into the bloodstream appeared in the subdural space, perhaps due to leaks from the vessels with fenestrated endothelium, but were stopped entering the subarachnoid space. In contrast, tracers injected into the CSF filled the SAS between the arachnoid and pia mater but did not cross into the subdural space. The authors claimed these tracers showed the existence of a single barrier beneath the dura mater—the arachnoid mater, and a single SAS. They further showed, using immunostaining, that the Prox1+ cell layer co-expressed VE-cadherin and clung to E-cadherin-positive ABCs, negating the possibility of CSF-filled space between them. Notably, they didn’t use Prox1-GFP transgenic mice, which could precisely show if the tracers were injected into the SAS beneath the Prox1+ cell layer. Hence, missing the Prox1+ layer in their *in vivo* optical imaging of the injected tracer may not be ruled out. Fluorescent VE-cadherin tagging could show the position of the arachnoid mater, but it couldn’t help precisely locate the barrier cell layer (Mapunda et al., 2023). Interestingly, the appropriate demonstration of arachnoid layers was also missing in Møllgård *et al*.’s *in vivo* optical imaging of the injected tracer in Prox1-GFP transgenic mice.

Using a reporter mouse for the ABC layer and Prox1 in tracer-based studies may be a way to examine the presence of SLYM as an independent leptomeningeal layer and compartmentalization of SAS.

Other than these studies, multiple e-letters in the original journal of publication (Møllgård *et al*., 2023) and comments on the science media platforms (*ALZFORUM*, 2023.) also questioned the existence of SLYM. The commenters generally believed that tracers appeared above the Prox1+ cells in Møllgård *et al*.’s experiments because the tracer was injected in the subdural space. Hence, they showed a subdural space created by the experimental manipulations, not a separate subarachnoid space.

The objections raised by the commenters seem rational, as Møllgård *et al*. didn’t use any marker to indicate the location of the ABC layer during optical imaging. Although the tracer they injected into the cisterna magna appeared beneath and didn’t cross through the Prox1+ layer, appropriate localization of the ABC layer has been a valid concern. The outer compartment they have shown seems to extend up to the dura mater. A missing arachnoid mater in optical imaging strengthens the view of commentators. Moreover, they also don’t precisely describe if the tracer was injected into the inner space beneath the Prox1+ layer of cisterna magna. Arguably, the compartmentalization of SAS by the SLYM layer should also extend into the cisterna magna.

In a recent article, Plá *et al*. presented further immunohistological evidence in postmortem mice brains to corroborate Møllgård *et al*.’s findings (Plá et al., 2023). Notably, both studies have a shared team of co-authors. Plá *et al*. precisely differentiated the SLYM from the arachnoid barrier cell (ABC) layer. The compartmentalization of SAS into the inner and outer spaces, including in the cisterns around the brain stem, was also demonstrated. However, in the mice’s spinal cord, the SLYM was shown to adhere to the ABC layer. Of note, the authors only examined the upper cervical segments of the spinal cord (Plá et al., 2023).

Plá *et al*. suggested that the collapse of the SAS during postmortem tissue processing may be a likely reason that the later studies that examined the existence of SLYM cannot show an intervening space between the arachnoid barrier and Prox1+ cells. The authors performed high-resolution (9 Tesla) serial MRI imaging of the postmortem mice brains to substantiate the SAS’s collapse during tissue processing (Plá et al., 2023).

However, neither Møllgård *et al*. or Plá *et al*. precisely demonstrated the trabeculae between the leptomeningeal layers. This fails to completely dispel the readers’ apprehension that the SAS compartments are not artifactually created due to the mechanical separation of the arachnoid cell layers during postmortem tissue processing. Alternative fixation methods that need shorter fixation time and immunostaining of fresh frozen sections may better preserve the SAS compartments and trabeculae between the leptomeningeal layers. Moreover, exploring the SLYM in larger animals, including humans, where SAS will be more voluminous, will help precisely differentiate it from the other leptomeningeal layers.

Surprisingly, Møllgård *et al*.*’s* and Plá *et al*. ‘s histological images show that the blood vessels are primarily limited to the inner space. It is very peculiar and anatomically challenging to explain, considering the established anatomy of the head-neck vessels. In the latter article, Plá *et al*. referred to the ultrastructural and light microscopic images from the earlier studies in the mammalian brain by Nabeshima *et al*. (Nabeshima et al., 1975) and Orlin *et al*. (Orlin et al., 1991) to support the compartmentalization of SAS by an intermediate leptomeningeal layer (Fig. 3).

**Figure 3.**
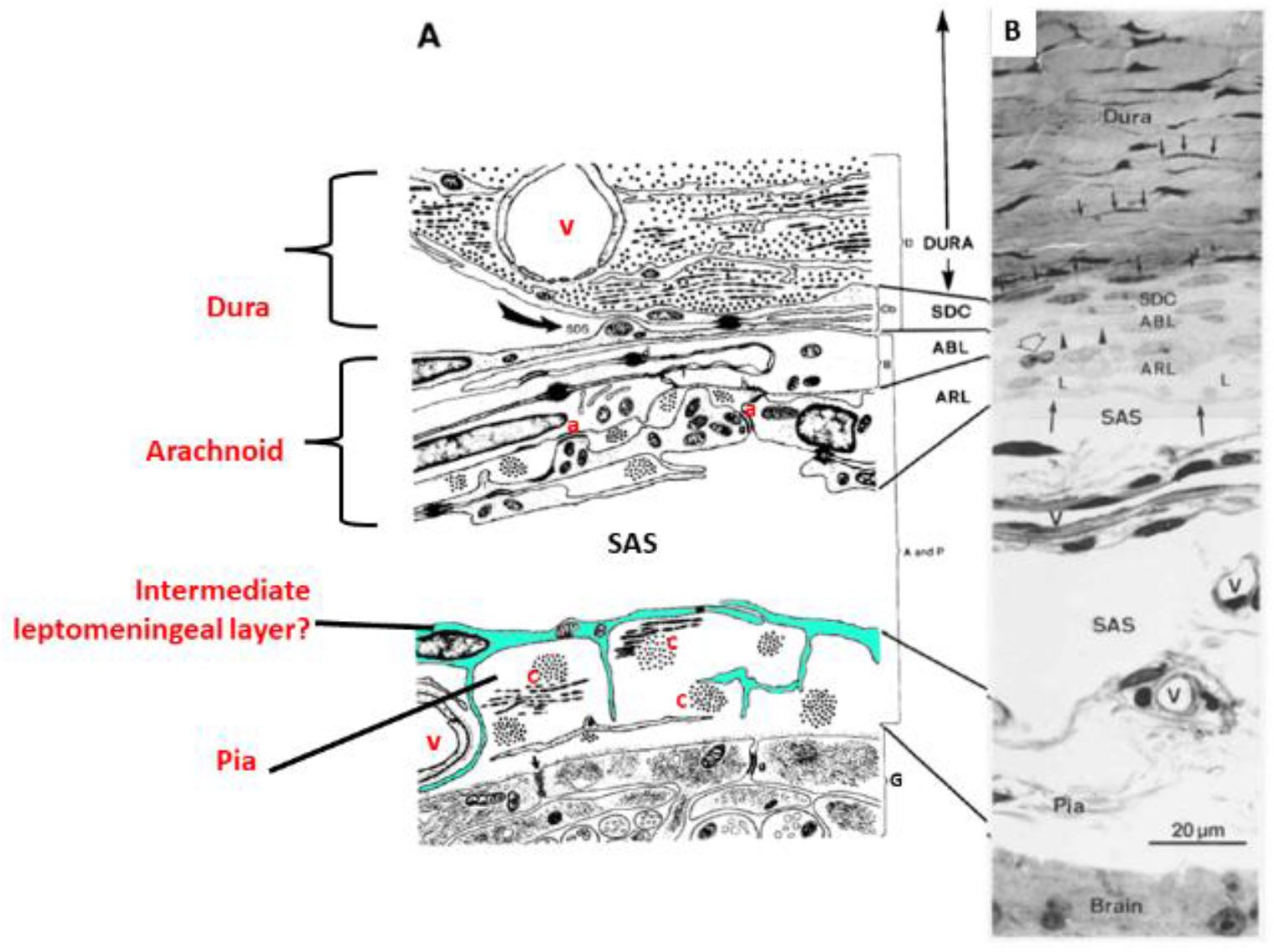
Schematic representation of the meningeal arrangement around the cerebral cortex originally demonstrated by A. Nabeshima et al., 1975 in TEM imaging of multiple mammals and B. Orlin et al., 1991 in light microscopic imaging of pigs. [A. A slightly modified schematic representation of meningeal arrangement around the cerebral cortex originally depicted by Nabeshima *et al*., 1975 based on his TEM-based results in multiple mammals. Original labelings are left in place, and any new labeling has been marked in red. A thick arachnoid mater is shown to have a dural border (Db) or subdural cell (SDC) layer, barrier (ABL), and reticular cell (ARL) layers. ABL is identified by characteristic tight junctions (t) and desmosomes (d) between the cells. The reticular cell layer is dominated by adherens junctions (a) and hemidesmosomes (h) and encloses cris-cross collagen bundles (C). A single-cell fibroblastic layer outer to the margin glia of the brain (G) represents the pia mater. An unexplained intermediate fibroblastic layer is distinctly observable between arachnoid and pia and is a likely candidate for the intermediate leptomeningeal layer. Junctional complexes between the cells are not well identifiable. Criss-cross collagen bundles (C) are enclosed between the intermediate layer and pia. B. A light microscopic image of thin toluidine sections of pig cortex by Orlin *et al*. 1991. His light microscopic image was highly comparable to the TEM-based schematic descriptions given by Nabeshima *et al*., showing a very similar arrangement of meningeal layers, including the intermediate fibroblastic layer between the arachnoid and pia mater. Small arrows: dural fibrocytes, arrowheads: dark intercellular line of the ABL, medium arrows: deep lining of the ARL, open arrow: a migrating cell. **Abbreviations:** A and P: Arachnoid-pia FBV: Fenestrated blood vessel in dura; B: Arachnoid barrier layer; G: Marginal glia of brain; C: collagen bundles; PVB: Pia-arachnoid blood vessel; D: Dura SAS: Subarachnoid space; Db: Dural border layer SDS: Subdural space; d: desmosomes; h: hemidesmosomes; t: tight junctions; a: adherens junctions; and g: gap junctions. **Reproduced with permission from Orlin *et al*., 1991**.]

Interestingly, the light microscopic image by Orlin *et al*., 1991 clearly shows the presence of blood vessels along the complete width of SAS. Also, the blood vessels in the annotated outer compartment in the image appear of a higher calibre than the inner compartment, substantiated by the narrower width of the latter (Fig. 3B). Due to its high evolutionary incentives, explaining whether the mice’s subarachnoid blood vessels have a different arrangement than the other mammals is crucial.

TEM-based validation of the SLYM and exploration of its detailed ultrastructure remains a concern. Although only a few TEM studies that examined this are currently available, none explicitly supported SLYM’s independent existence (Grubb, 2023; Mapunda et al., 2023; Pietilä et al., 2023). Grubb recently analyzed publicly available electron microscopy datasets for mice brains (Grubb, 2023). The author described the arachnoid mater as composed of at least four different cell types: the dural border cells, ABCs, reticular cells, and inner arachnoid fibroblasts. The arachnoid mater comprises an inner layer of fibroblasts that attach to and are indistinguishable from pia mater fibroblasts and resemble the BFB2 cells identified by Pietilä *et al*. Above that is a layer of reticular cells, which resemble the BFB3 fibroblasts identified by Pietilä et al. The author speculated that the reticular cell layer may be the SLYM reported by Møllgård et al. The author further noted that reticular cells are fenestrated and are loosely attached among themselves and to ABCs with adherence- and gap junctions. They further stated that the reticular cells have a large amount of rough endoplasmic reticulum and large mitochondria. They often have more than one nucleus and appear darker (more electron-dense) than the other cells in the arachnoid mater. Outer to the reticular cell layer lies the ABC layer. These cells are often multinuclear (up to 4 nuclei), form a tight barrier, and are attached by long stretches of adherence and gap junctions with slim and electron-dense intracellular spaces (Grubb, 2023). The author opined that the inter-cellular junctions of the reticular cell layer (with each other and with the ABC layer) could be VE-cadherin instead of E-cadherin, as indicated in the study by Mapunda *et al*. (Grubb, 2023; Mapunda et al., 2023). Grubb’s view is supported by all contesting studies, which showed E-cadherin located explicitly in the ABC layer (Grubb, 2023; Mapunda et al., 2023; Møllgård et al., 2023; Pietilä et al., 2023).

The possible reasons for the variance of opinion about the intermediate leptomeningeal layer in TEM studies must be resolved. Notably, TEM imaging requires a tiny sample. In little brain samples, leptomeningeal layers may get lost during tissue processing, thus confounding the interpretation of the ultrastructural images. The other factors, such as postmortem delay, type of fixative, and fixation duration, may also impact the salvage of the cellular layers during histological processing for TEM. Moreover, the intermediate leptomeningeal layer may have an irregular placement across the SAS; hence, in ultrastructural imaging, it may be confused with the obliquely placed arachnoid trabeculae. Notably, horizontally placed trabecular sheets were described in the previous ultrastructural studies.

### 4.4 Hidden indications in published SEM data

A comprehensive review of the prior dissection-based or microscopic studies in animal/human anatomy gives no clear indication of the presence of SLYM. Despite being a macroscopic structure, how it escaped from the observations in the scanning electron microscopic (SEM) studies is astonishing. As a limited stance, Alcolado *et al*. in 1988 indicated the existence of sheet-like trabeculae in the SAS of the brain, which consisted of collagen bundles and leptomeningeal cells connected by desmosomes and gap junctions (Alcolado et al., 1988). However, they gave no indications of any intermediate leptomeningeal layer. In the same year, Nicholas and Weller described the presence of an intermediate meningeal layer between the arachnoid and pia mater in the human spinal cord using SEM (Nicholas & Weller, 1988). Albeit they did not demonstrate its presence in the brain. Unfortunately, their discovery was not adequately followed up in later studies.

A review of the published data hints that a miss of the SLYM in the SEM could have been caused by an error of annotations in the absence of factual knowledge about this layer. Below, we discuss the SEM images of CNS components from the published literature in mammals (Saboori, 2021; Saboori & Sadegh, 2015; Roy O. Weller et al., 2018). Figs. 4-5 show the structural components of the SAS in the brain, Fig. 6 in the spinal cord, and Fig. 7 around the optic nerve. In Fig. 4, the membranous structure annotated as the pia mater by the original authors could be the new meningeal layer, as the SAS can be observed on both sides of this structure. The pia mater is visible as the membrane firmly adhered to the brain tissue. The intermediate layer is seen to give attachment to the trabeculae (more evident in the side of the arachnoid mater). This layer is much thinner than the arachnoid mater but thicker than the pia mater and appears to have a fibrous component. The SAS is well visible on both aspects of the intermediate layer.

**Figure 4.**
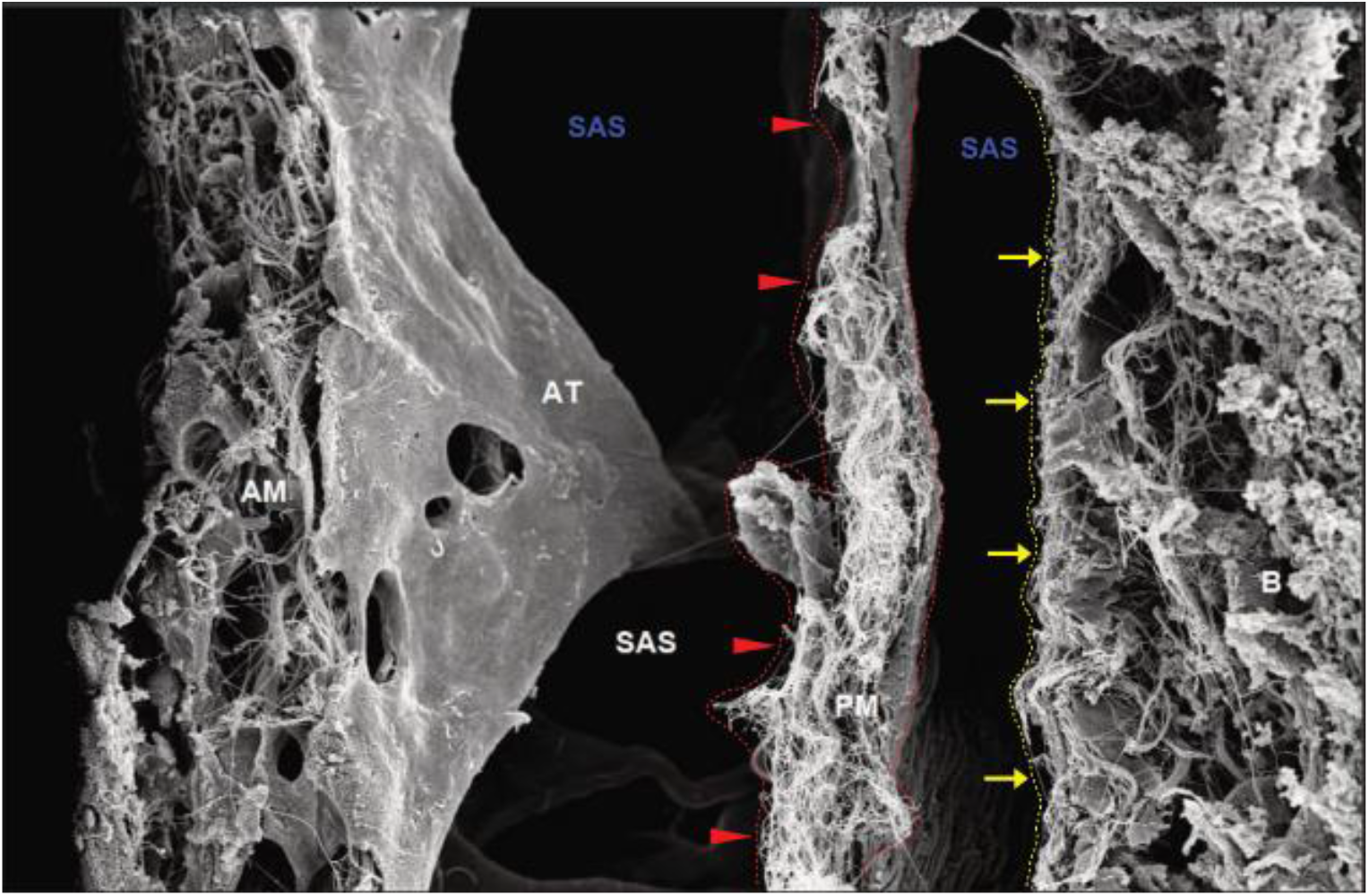
A scanning electron microscopic view of the subarachnoid space in a rat brain. (Magnification 2500 X, Scale bar 10 μm) [The arachnoid mater (AM) and arachnoid trabeculae (AT) are well marked. The pia mater is visible as the membrane adhered to the brain tissue (shown with yellow arrows). The membranous structure originally annotated as the pia mater (PM) by Saboori (2021) (shown with red arrowheads) is likely to be the new meningeal layer, as the subarachnoid space (SAS) can be observed in both aspects. This intermediate layer is seen to give attachment to the trabeculae (more evident in the side of the arachnoid mater). [The fluorescent color marks indicate re-annotations in the original image. **Reproduced with permission from Saboori, 2021**.]

**Figure 5.**
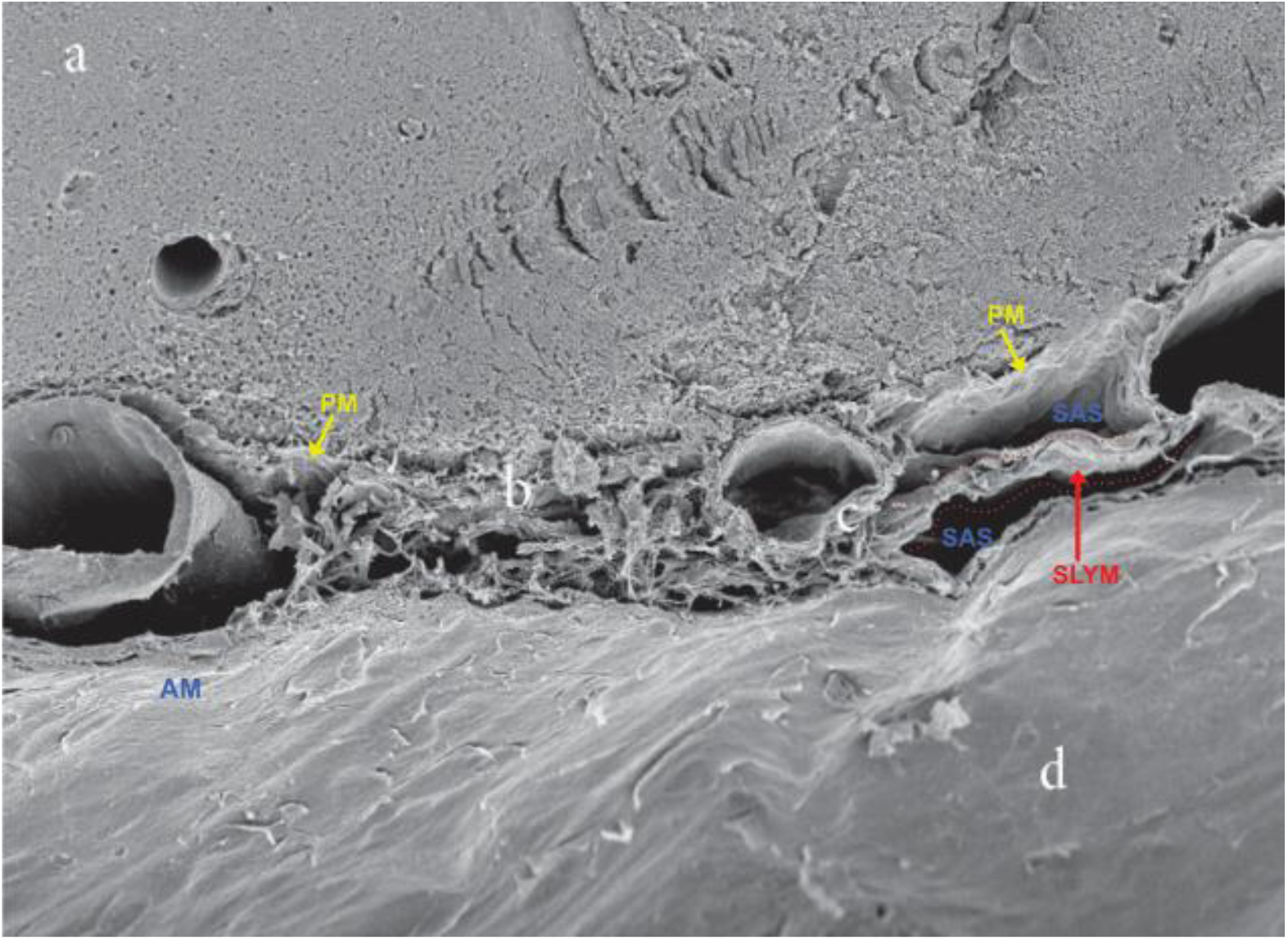
A scanning electron microscopic view of the subarachnoid space in a rat brain. [The dura mater (d), arachnoid mater (AM), trabecular network (b), and pia mater (PM) are marked. The blood vessels (c) can be seen ensheathed with a membranous layer (shown with red dotted lines), likely to be the new meningeal layer. This membranous layer divides the subarachnoid space (SAS) into two compartments; however, this is not traceable in the complete length of the section. [The fluorescent color marks indicate re-annotations in the original image. **Reproduced with permission from Saboori and Sadegh, 2015**.]

**Figure 6.**
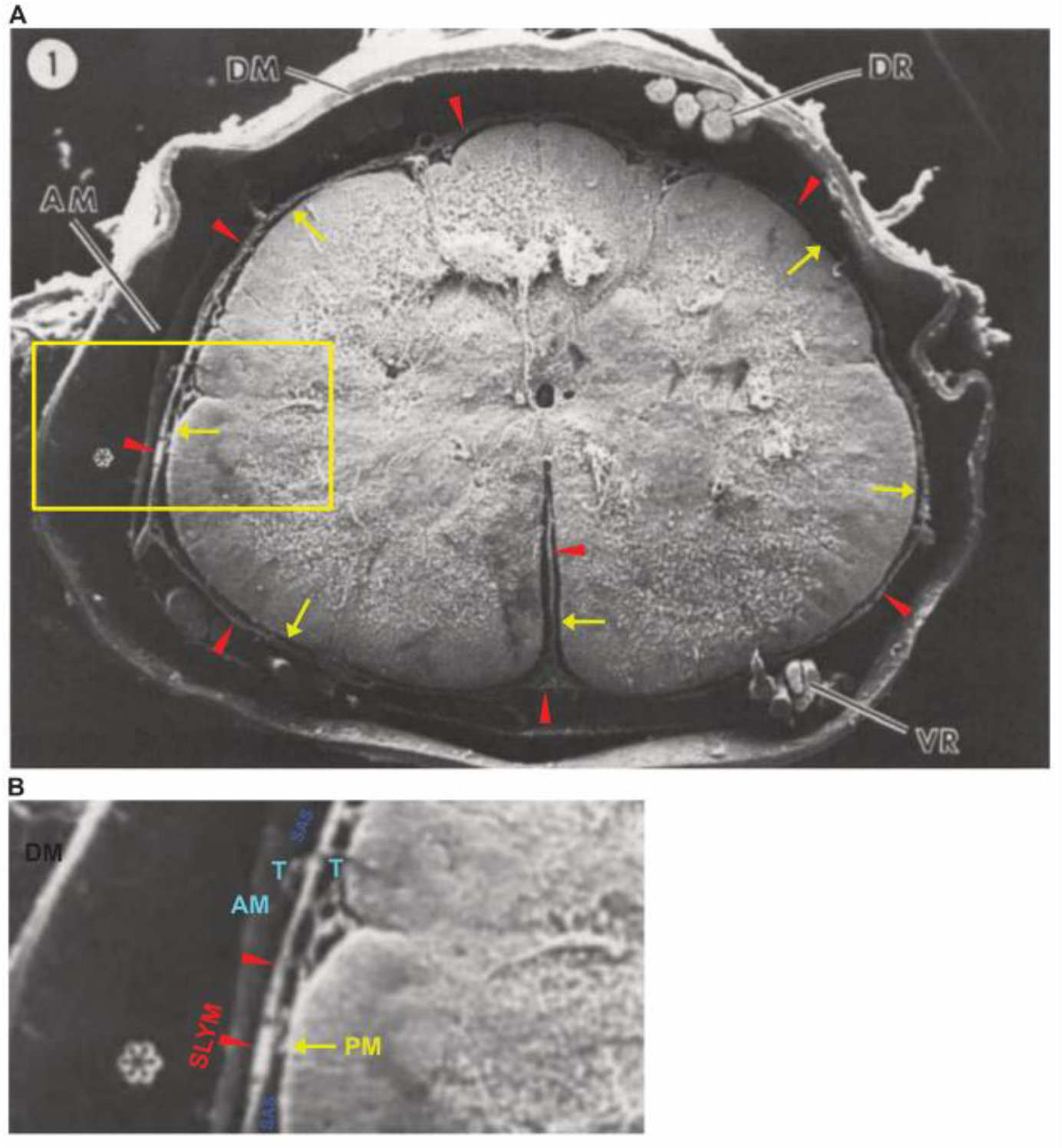
A scanning electron microscopy of the cross-section of a dog’s spinal cord and meninges. [A. The dura mater (DM) and arachnoid mater (AM) are well marked. The asterisk indicates the subdural space. The pia mater (PM) closely invests in the neural tissue and dips into the fissures (marked with yellow arrows). An intermediate membranous layer between the arachnoid and pia is likely SLYM: the new meningeal layer (marked with red arrowheads). This layer is loosely apposed to the neural surface and passes over the fissures. It sends a facial septum into the depth of the anterior median fissure. B. An Inset image. CSF-filled spaces and trabeculae (T) can be observed on both aspects of the intermediate layer. Abbreviations: SLYM: subarachnoid lymphatic-like membrane, SAS: subarachnoid space. [The fluorescent color marks indicate re-annotations in the original image. Reproduced with permission from Merchant and Low, 1979.]

**Figure 7.**
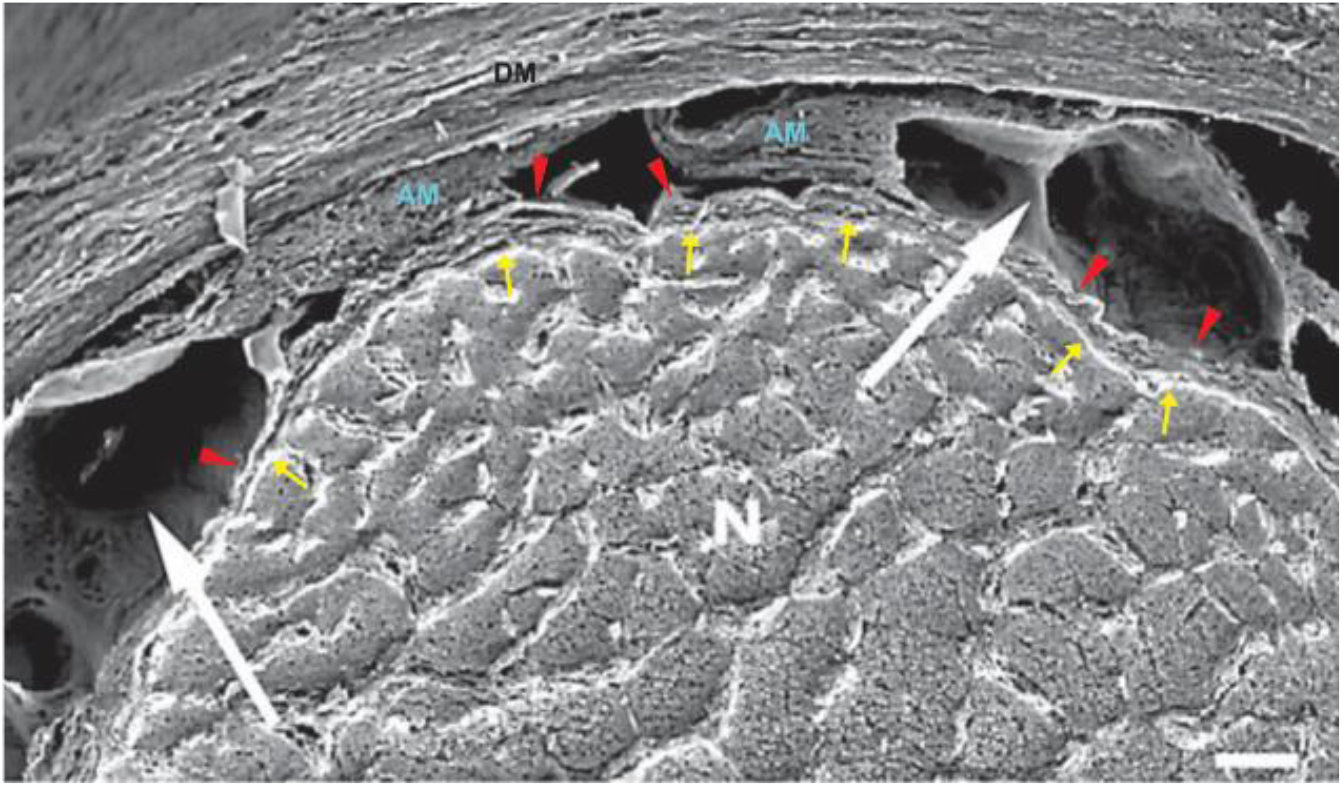
A scanning electron microscopic view of the subarachnoid space around the human optic nerve. [The dura mater (DM) and arachnoid mater (AM) are marked. The pia mater is indicated by a thin white sheath encircling the nerve (shown with yellow arrows). The membranous layer shown with red arrowheads is likely the new meningeal layer. This membranous layer divides the subarachnoid space (SAS) into two compartments. The outer compartment contains prominent trabeculae and septae (shown with white arrows). The inner compartment is narrow, but the delicate trabeculae forming a meshwork are observable. (The fluorescent color marks indicate re-annotations in the original image. **Reproduced with permission from H E Killer et al., 2003**.]

Similarly, in Fig. 5, an intermediate layer is observable in a part of the SAS. However, at other places, it seems confounded by the presence of arachnoid trabeculae. The blood vessels seem to be chiefly present in the inner compartment of SAS, limited by the intermediate layer and pia mater.

Fig. 6 presents a reannotation of a published SEM image of the cross-section of the dog’s spinal cord and covering meninges (Merchant & Low, 1979). Our re-annotation indicates the pia mater is firmly adherent to the spinal cord and dipped into the spinal fissures (Fig. 6A, indicated by yellow arrows). There is another membranous structure underneath the arachnoid mater. This is likely the new meningeal layer, free from neural surfaces and passing over the spinal cord fissures. It divides the SAS into two compartments (Fig. 6A, indicated by red arrowheads). The presence of CSF-filled spaces and trabeculae can be marked in both aspects (Fig. 6B, Inset image). Interestingly, this membranous structure encloses the anterior spinal vessels, sending a septum into the anterior median fissure (Fig. 6A, indicated by red arrowheads).

In Fig. 7, the membranous structure shown with red arrowheads is likely the new meningeal layer. This membranous layer divides the subarachnoid space (SAS) into two compartments. The outer compartment is visible and contains prominent trabeculae and septae (shown with white arrows). The inner compartment is not so obvious; however, a fine trabecular meshwork is observable inside it. Although the original authors described this layer as the pia mater (Killer et al., 2003), it shouldn’t bear a trabeculated space beneath to be considered that. In our view, the actual pia mater is indicated by a thin white sheath closely encircling the nerve (shown with yellow arrows).

These observations suggest a revised interpretation of the SEM images of SAS in light of the demonstration of the intermediate leptomeningeal layer in the brain by Møllgård *et al*. (Møllgård et al., 2023). Apart from the lack of factual information, there could be multiple reasons why the SEM studies missed this macroscopic structure. A thin membrane lacking robust fibrous support could be at a high risk of getting damaged during tissue processing. Moreover, its identification could be confused because it is sandwiched between arachnoid trabeculae and possibly its uneven placement in the collapsed SAS. Specifically, the flattened and obliquely placed trabeculae can confound its identification. As its presence in the brain is established, it can be more explicitly targeted in SEM imaging. Following it along the complete length of SAS may help differentiate it from trabeculae.

### 4.5 Possible regional differences in SLYM structure

Møllgård *et al*.’s description of SLYM in mice and humans is limited to only the cortex and brain stem. The SLYM has not yet been shown in the spinal cord or around the optic nerve. Both structures are covered with the meninges, showing the continuity of SAS with the brain. The rare description of the epipial tissue layer (Millen & Woollam, 1961) or intermediate leptomeningeal layer (Nicholas & Weller, 1988) between arachnoid and pia mater in the human spinal cord varies significantly from that by Møllgård *et al*. (Møllgård et al., 2023) in the brain. Millen and Woollam (Millen & Woollam, 1961), as well as Nicholas and Weller, described a fenestrated membrane that can allow the CSF content exchange (Nicholas & Weller, 1988) (Fig. 8).

**Figure 8.**
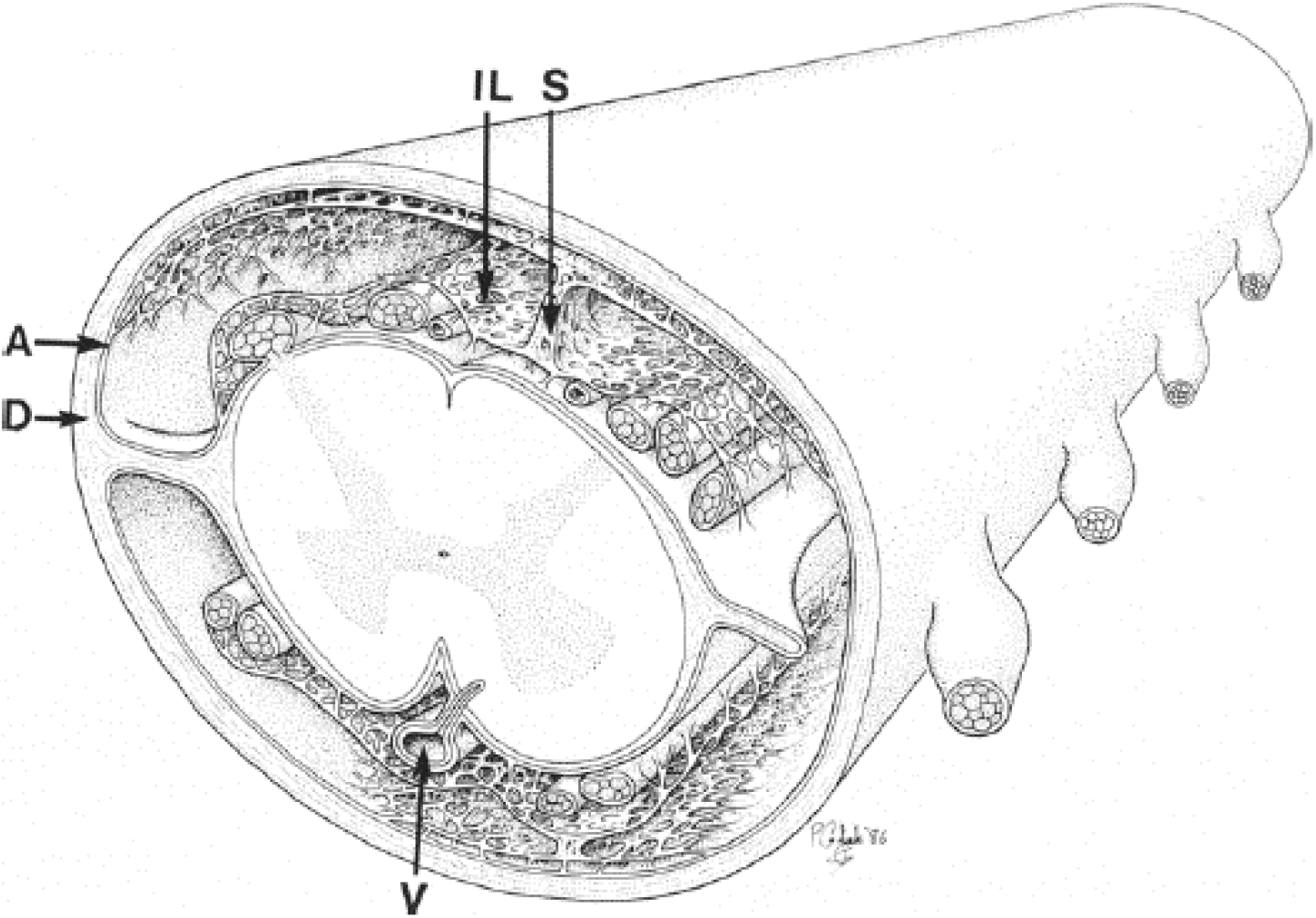
The diagrammatic representation of the meningeal arrangement in the human spinal cord by Nicholas and Weller, 1988. [Nicholas and Weller, as early as 1988 (Nicholas & Weller, 1988) described the presence of an intermediate leptomeningeal layer (IL) between the arachnoid and pia mater. The IL is loosely attached to the inner aspect of the arachnoid (seen closely applied to the dura) but is distinct from it and reflected in the midline to form dorsal septa that traverse subarachnoid space. Its fenestrated lateral portions formed delicate filiform trabeculae connected to the pia, which enclosed blood vessels and nerve roots. They further showed that it was possible to see the underlying pia mater by peering through the fenestrations in the IL (seen covering the spinal cord). IL was less substantial than seen dorsally in the ventral aspect of the spinal cord, and no ventral septum was formed. **Abbreviations**: IL: Intermediate Leptomeningeal layer; D: Dura; A: arachnoid; S: Septa; V: Vessel. **Reproduced with permission from Weller et al**., **1992.]**

In contrast, Møllgård *et al*. showed that SLYM is an impermeable structure that does not allow the passage of moieties more than one μm and three kilodaltons (Møllgård et al., 2023).

Although the studies by Millen and Woollam (Millen & Woollam, 1961), Nicholas and Weller (Nicholas & Weller, 1988), and Møllgård *et al*. (Møllgård et al., 2023) used different methods, the mismatch of the conclusions is evident. This hints at possible regional differences in the architecture of the SLYM. It will be necessary to investigate its architecture along the SAS’s complete length, especially along the spinal cord and optic nerve, to understand its impact on CSF dynamics and brain barrier functions.

### 4.6 Hidden Indications in Neuroimaging

Among the commonly practiced neuroimaging methods, ultrasonography (USG) and computed tomography (CT) scans are mainly limited to detecting the dura mater. Magnetic resonance imaging (MRI), the most sensitive imaging modality for viewing the meninges, can see all three meningeal layers (Cinnamon et al., 1994; Kirmi et al., 2009; Meltzer et al., 1996; Patel & Kirmi, 2009; Sze, 1993; Trinh & Massoud, 2023). The leptomeningeal layers, specifically the pia mater, may be complex to appreciate due to the thinness. The leptomeningeal layers show an enhancement in MRI owing to their high vascularity, mainly when contrast is used. The T2-weighted MRI can be more suitable for viewing the leptomeningeal layers as they appear hypointense compared with hyperintense CSF sandwiched between them (Cinnamon et al., 1994; Kirmi et al., 2009; Meltzer et al., 1996; Patel & Kirmi, 2009; Sze, 1993; Trinh & Massoud, 2023). The arachnoid mater and pia mater are more clearly differentiated at the sulci, fissures, and cisterns owing to the increased gap between the layers. The SAS gets deeper at these locations with ample CSF in between, which can help differentiate the leptomeningeal layers (Standring, 2021).

Analyzing a typical MRI image from an adult human brain does not indicate an intermediate structure dividing SAS. Using the T2 weighted or FLAIR, or contrast-enhanced images, or applying the ultra-high magnetic field, such as 7 Tesla, seems to provide no help. However, a thin hypointense line can be seen traversing longitudinally through the middle of SAS in T2 weighted images of the spinal cord (Fig. 9). This line is observed bilaterally in the complete length of SAS and hints at a fascial structure (Fig. 9). Further, the line can be traced upward over the cisterna magna (Fig. 9). Being a continuous line, and the presence of CSF on both aspects indicate it to be a membranous structure, most likely the SLYM. The non-visibility of any such streak in the brain SAS in T2 weighted MRI images of the normal brain can be explained by the narrow width of the SAS.

**Figure 9.**
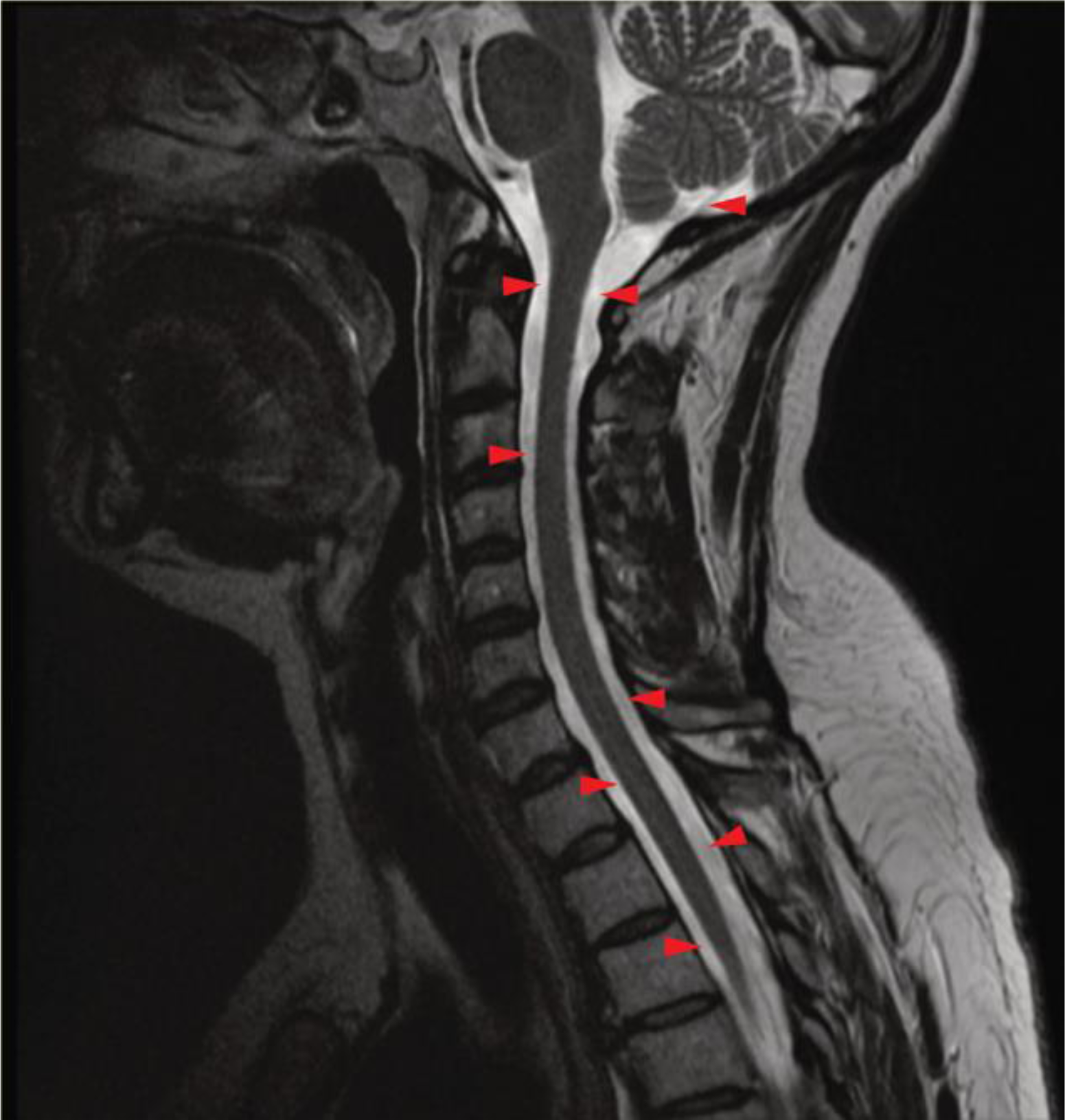
Saggital section of T2 weighted MRI of an adult human spine. A thin hypointense line traverses longitudinally through the middle of SAS bilaterally in the spinal cord (marked by red arrowheads) in its complete length. It can be traced upward over the cisterna magna. The continuity of this line in the entire length of spinal SAS and the presence of hyperintense CSF on both aspects indicate this could be a facial layer, perhaps SLYM (**Image source: Case courtesy of Frank Gaillard, Radiopaedia.org, rID: 35630, http://creativecommons.org/licenses/by-nc/4.0/**, (Gaillard, 2015)).

This view is supported by the faint visibility of an intermediate hypointense line in the T2 weighted brain MRI images of the cases of benign enlargement of SAS (BESS) in infants (Fig. 10). A wide SAS may help to differentiate the identification of the intermediate hypointense structure against the backdrop of hyperintense CSF. This middle hypointense line in SAS in BESS cases can be seen in the full extent of the brain (Fig. 10).

**Figure 10.**
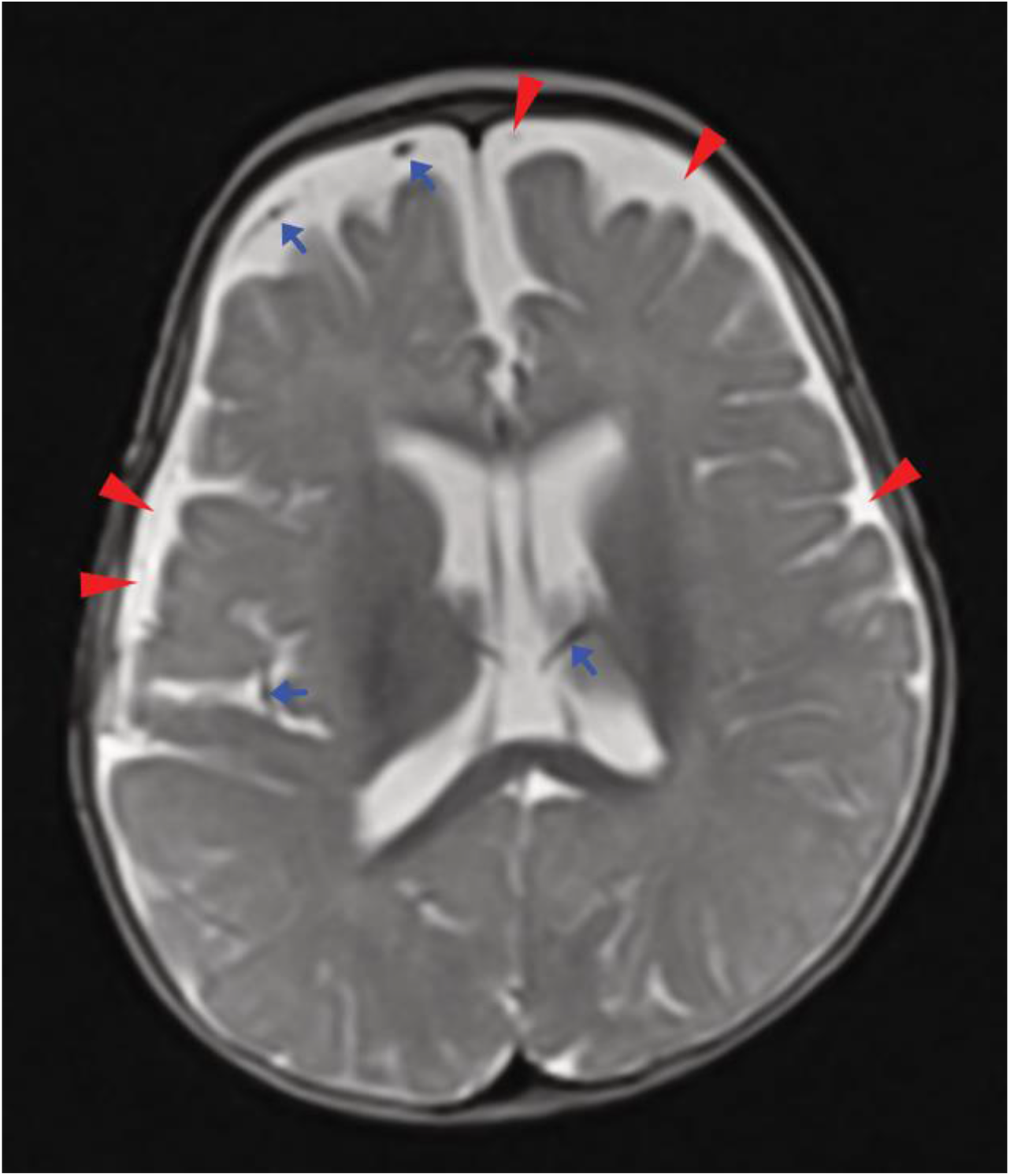
Axial section of T2 weighted MRI in a case of benign enlargement of SAS in infants. [The wide SAS helps to differentiate an intermediate hypointense line against the background of hyperintense CSF (marked by red arrowheads). This line passes over the sulci and is seen to the full extent of the brain, perhaps a membranous structure representing SLYM. The blood vessels can be differentiated from SLYM, being more hypointense and wide solid lines (marked by blue arrows). (**Image source: Case courtesy of Heba Abdelmonem, Radiopaedia.org, rID: 59619, http://creativecommons.org/licenses/by-nc/4.0/**, (Heba, 2018.)]

Optical coherence tomography (OCT), an optical imaging modality used for high resolution cross-sectional imaging can potentially visualize SAS components (Benko et al., 2020; Hartmann et al., 2019). It can be used *in vivo* in the neurosurgery patients who go for craniotomy or *in situ* in postmortem brains after opening the cranial volt. We re-annotate a published OCT image of SAS in fresh frozen cadaveric human brains, indicating its plausible compartmentalization by SLYM (Fig. 11) (Benko et al., 2020).

**Figure 11.**
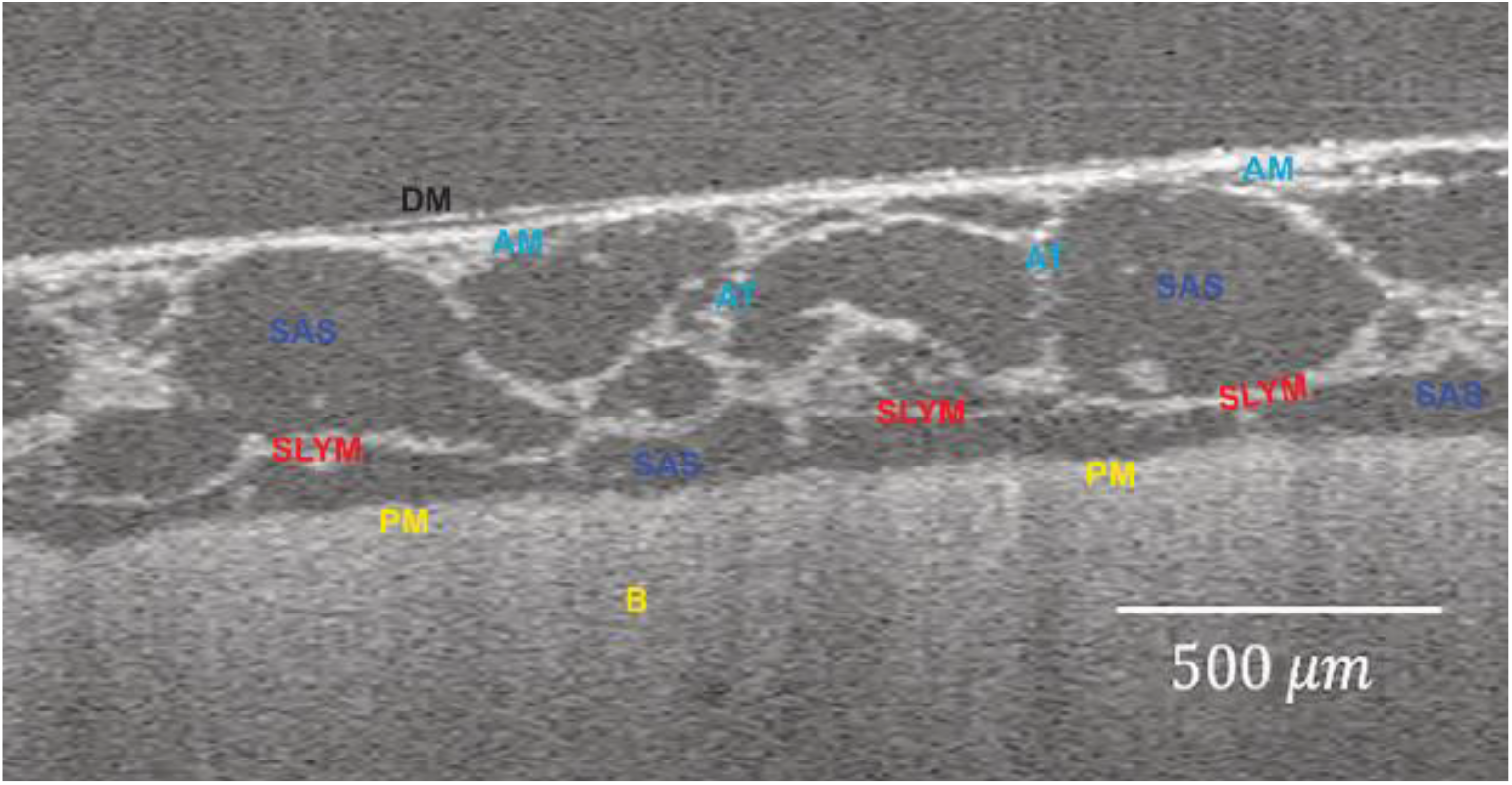
In situ optical coherence tomography imaging of the subarachnoid space in postmortem human brain. [The dura mater (DM), arachnoid mater (AM), and pia mater (PM) are marked. The subarachnoid space (SAS) is traversed by arachnoid trabeculae (AT). A transverse septum is seen all along the brain (B) surface and divides the SAS into two compartments (marked as SLYM), which can be the new meningeal layer. The fluorescent color marks indicate re-annotations in the original image. **Reproduced with permission from Benko et al**., **2020**.]

### 4.7 Physiological and clinical relevance

Functional compartmentalization of the SAS by SLYM necessitates a new conceptualization of the dynamic of CSF circulation. Per the traditional description, the CSF is secreted through the choroid plexus in the cerebral ventricles (chiefly in the lateral but also the third and fourth). From the lateral ventricle, CSF enters the third ventricle through the interventricular foramen (of Monro) and then through the cerebral aqueduct into the fourth ventricle. From the fourth ventricle, it enters into SAS around the brain and spinal cord through the openings in the inferior aspect of the roof of the fourth ventricle (foramen of Magendie in the midline and foramen of Luschka bilaterally). From SAS, CSF is drained into the dural venous sinuses through arachnoid granulations (Standring, 2021). The presence of the SLYM has now complicated the earlier described course of CSF through SAS. It is currently not known if the outer and inner compartments created by the SLYM have any anatomical communication along the complete length of SAS. The compartmentalization may have an impact on the composition of the CSF. It is also unknown whether CSF pressure and CSF circulation vary between the compartments. Arguably, the inner compartment bounded between the new membrane and pia mater lies in a close approximation of the brain parenchyma; it will exclusively receive the brain waste.

On the contrary, the outer compartment will contain a clearer CSF, as there is a limited exchange of solutes between the two compartments. The CSF has an established role in clearing brain waste (Plog & Nedergaard, 2018). The presence of SLYM creates an additional layer in the blood-CSF/brain barrier. Moreover, the blood vessels passing through the inner compartment may carry a sleeve of SLYM; hence, it is also likely to contribute to the glymphatic system (Møllgård et al., 2023), the proposed perivascular route of waste clearance from the CNS (Kumar et al., 2019; Plog & Nedergaard, 2018).

The compartmentalization of SAS by SLYM complicates the course, how the CSF exiting through the median and lateral openings at the fourth ventricle enters into two separate compartments, and how it is absorbed into the dural venous sinuses. As SLYM does not allow passage of molecules greater than one μm and three kilodaltons (Møllgård et al., 2023), it is unclear how larger molecules generated as brain waste will be removed from the inner compartment if it has no direct access to the dural venous sinuses. Alternatively, the glymphatic route is the most likely explanation for the clearance of the solute waste from the inner compartment.

Contrary to the belief that the brain is an immune-privileged organ, the physiological presence of immune cells, primarily leucocytes, in meningeal spaces and circulation in CSF is now an established phenomenon (Gadani et al., 2017; Nevalainen et al., 2022; Walker et al., 2018; Wolf et al., 2009). The meningeal immune cells are believed to monitor the entry of pathogens to the CNS. Møllgård *et al*. showed that SLYM not only shares the immunophenotypic characteristics with the lymphatic vessels but also shows the presence of receptors of immune cells (Møllgård et al., 2023), which indicates that it creates an immunogenic checkpoint against the pathogens invading the SAS. A traumatic rupture or pathological degeneration of SLYM will allow the mixing of the dirty and clear CSF and permit the access of pathogens in the inner compartment and, consequently, to the CNS parenchyma. Thus, an intact and healthy SLYM creates an additional protective barrier to the CNS.

Aging and consequent immune cell dysregulation have been implicated in the pathogenesis of many neurodegenerative disorders, such as Alzheimer’s and Parkinson’s, amyotrophic lateral sclerosis, and multiple sclerosis (Mayne et al., 2020). Possible degenerative changes in the SLYM with aging may lead to their pathogenesis.

The discovery of the new meningeal layer also necessitates revising the drug delivery approaches to the CNS. Inter-individual variations in the clinical and adverse effects of subarachnoid drug administration for spinal anesthesia/analgesia are common (Fettes et al., 2009). The anatomical factors are suggested to play a role (Tuominen et al., 1992); however, the exact mechanisms have never been understood. The SLYM can now help to explain this phenomenon. The neural absorption of an anesthetic agent injected into the respective CSF compartments of spinal SAS may vary, which will be reflected in their clinical and adverse effects.

### 4.8 Limitations and future directions

Møllgård *et al*.’s SLYM description is limited to its identification as a new meningeal layer (Møllgård et al., 2023). Very little is known about its structure, especially macroscopic appearance and ultrastructural details. The controversy around its existence in recent studies (Mapunda et al., 2023; Pietilä et al., 2023) must be resolved.

Although SLYM has to be a macroscopically visible structure, approaching it in the animal or human brain remains challenging, limiting its neurosurgical application and anatomical demonstration during medical teaching. Further, its topographical arrangement around the brain surfaces—particularly at sulci and cisterns, vascular relations, contribution to the blood-

CSF/brain barriers, and CSF circulation, including secretion and absorption through the compartments, are currently least understood. Also, the CSF contents in the outer and inner compartments will likely vary. Moreover, no other significant purpose of the compartmentalization of SAS is clear except for creating an immunogenic barrier. These aspects must be extensively studied to understand SLYM’s complete physiological and clinical significance.

The presence of the SLYM also necessitates a rethinking of the anatomical recognition of some meningeal structures which are considered modifications of the pia mater, such as tela choroidea in the brain, and *ligamentum denticulatum, linea splendens*, and *filum terminale* in the spinal cord. These meningeal modifications are seen just beneath the arachnoid mater; hence, keeping with traditional description, it is likely that their identification might have been erroneously imparted to the pia mater (Mizutani & Rodesch, 2020).

Despite frequent previous mentions of the intermediate leptomeningeal layer in brain and spinal SAS, it has been little appreciated in the current literature. Available descriptions in the literature are inconsistent and use varying names, which can explain this obscurity. Its thorough exploration along the complete length of the central nervous system will be desirable to establish it as an independent meningeal layer. Moreover, devising the neurosurgical approaches for cadaveric dissection and *in vivo* operative procedures and using standard terminology in line with the existing meningeal layers will be essential to popularise the newly discovered meningeal layer in medical education and clinical practice.

### 4.9 Concluding remarks

Occasional studies described the presence of an intermediate meningeal layer in the SAS of the brain or spinal cord, which closely resembled the newly discovered meningeal layer. However, unequivocal descriptions for this layer all along the central nervous system are scarce. Its detection in neuroanatomical and imaging-based studies, especially in the brain, could have been missed for various reasons, such as extreme thinness, loss during tissue processing, and confounding with the arachnoid trabeculae and blood vessels in the SAS. The discovery of the SLYM is a landmark addition to the neuroanatomy and will likely redefine it in multiple aspects. It is expected to change the existing concepts about the protective barriers and the dynamic circulation of CSF around the CNS.

Consequently, established conceptions about the pathogenesis of many neurodegenerative diseases are also likely to change. It can potentially also revolutionize the drug delivery approaches to the CNS. Its discovery seems to open a new avenue in neuroscience research, and future studies are likely to unravel its detailed structure and role in brain health and disease.

Although prior descriptions in literature corroborate SLYM, controversy persists about its independent existence. A potential caveat with SLYM discovery is that it has not yet been established in ultrastructural studies. Its detailed structure and function also remain to be established. Future studies may hopefully fill these gaps.

## Acknowledgment

The authors express their heartfelt gratitude to Professor Parisa Saboori, Manhattan College, Riverdale, NY, USA, for permitting the reuse/adaptation of figures from her published articles.

## Author (s) contributions

A.K. designed and A.K., C.K., and A.A. supervised the study. A.K., R.K., R.K.N., R.K.J, and A.A. collected and analyzed data. A.K., R.K., R.K.N, B.N. prepared figures and tables. A.K., R.K., and R.K.N wrote the first draft. C.K., A.A., P.K., A.K.R., A.K.D, V.P., P.P., M.A.F, S.N.P, and P.A. reviewed for, and all authors consented to the final draft.

## Data availability statement

Data sharing does not apply to this article as no datasets were generated or analyzed during the current study.

## Supplementary files

**Supplementary file 1, Search strings**.

(“meninges”) AND layers AND (subarachnoid space OR SAS)

(“meninges”) AND layers AND brain AND (subarachnoid space OR SAS)

(“meninges”) AND layers AND brain AND inflammation AND immune cells (subarachnoid space OR SAS)

(“meninges”) AND layers AND spinal cord AND (subarachnoid space OR SAS)

(“meninges”) AND development* AND mesothelium AND (embryonic*)

(“meninges” OR “dura mater” OR “arachnoid” OR “pia mater”) AND development* AND (embryonic*)

(“meninges”) AND layers AND central nervous system AND (subarachnoid space OR SAS)

(“meninges”) AND microscopy AND electron AND scanning AND (subarachnoid space OR SAS)

(“arachnoid”) AND microscopy AND electron AND scanning AND (subarachnoid space OR SAS)

(“pia mater”) AND microscopy AND electron AND scanning AND (subarachnoid space OR SAS)

(“epipial layer” OR “inner pial layer” OR “outer pial layer”) AND (brain OR spinal cord)

(“meninges”) AND microscopy AND electron AND transmission AND (subarachnoid space OR SAS)

(“meninges”) AND magnetic resonance imaging AND (subarachnoid space OR SAS)

(“meninges”) AND ultrasonography AND (brain OR spinal cord) AND (subarachnoid space OR SAS)

(“meninges”) AND computed tomography AND (brain OR spinal cord) AND (subarachnoid space OR SAS)

(“meninges”) AND anatomy* AND (subarachnoid space OR SAS) AND (humans OR primates or rats or mice or animals)

(“meninges”) AND blood-brain barrier AND BBB AND (subarachnoid space OR SAS)

(“meninges”) AND blood-CSF barrier AND (subarachnoid space OR SAS OR trabeculae)

(“meninges”) AND cerebrospinal fluid AND circulation AND (subarachnoid space OR SAS)

(“meninges”) AND layers AND spinal cord AND (ligamentum denticulatum OR linea splendens OR filum terminale)

(“meninges”) AND layers AND brain AND humans AND (dissection OR anatomy)

(“meninges”) AND layers AND brain AND inflammation OR immune cells (subarachnoid space OR SAS)

(“meninges”) AND layers AND brain AND lymphatic vessels AND (subarachnoid space OR SAS)

(“meninges”) AND layers AND brain AND glymphatic system AND (subarachnoid space OR SAS)

(“meninges”) AND layers AND (subarachnoid space OR SAS) AND subarachnoid lymphatic-like membrane OR SLYM

(“meninges”) AND central nervous system AND clinical significance AND (subarachnoid space OR SAS)

**Supplementary file 2,**

**Figure S1.**
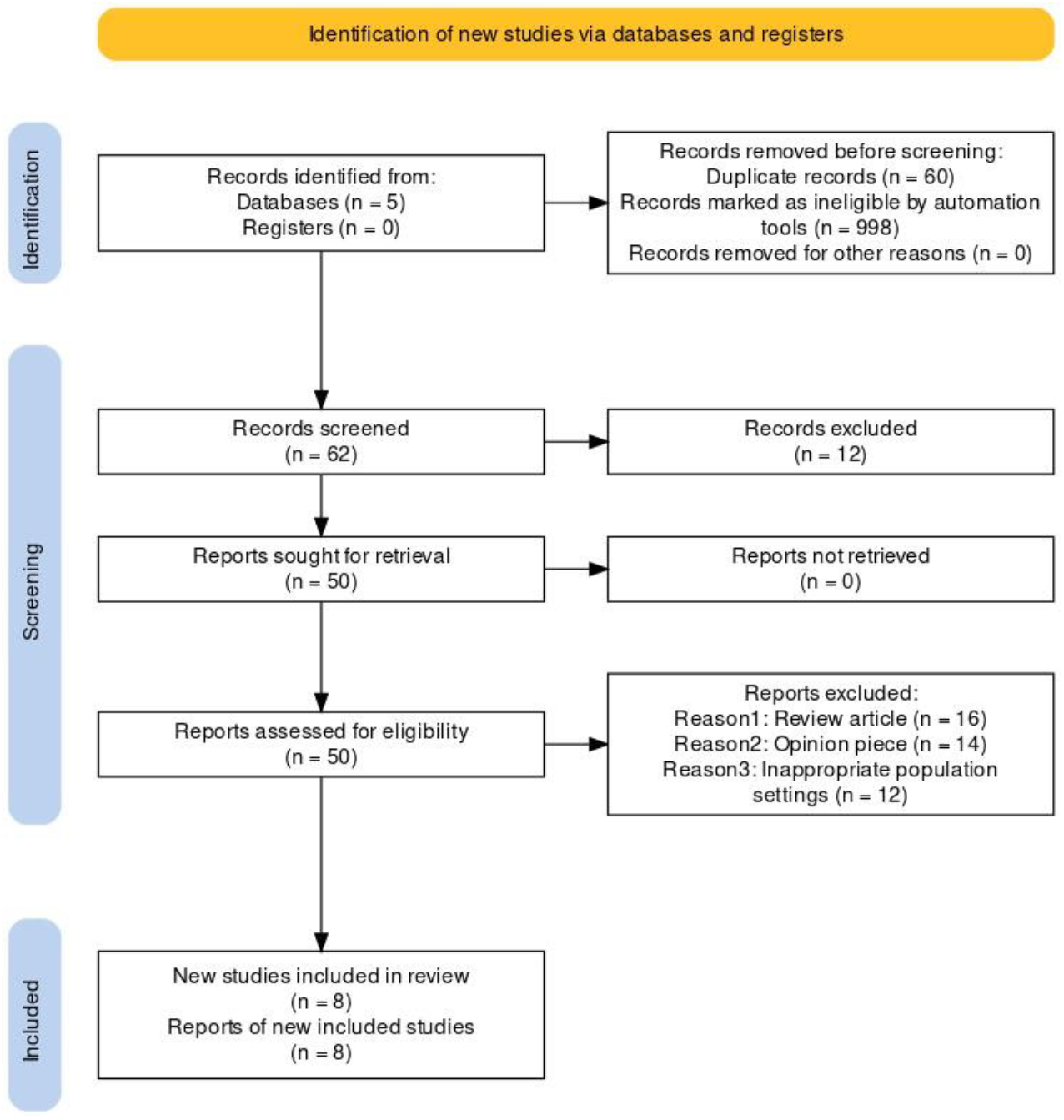
PRISMA2020 flow diagram.

**Supplementary file 3,**

**Table S1.**
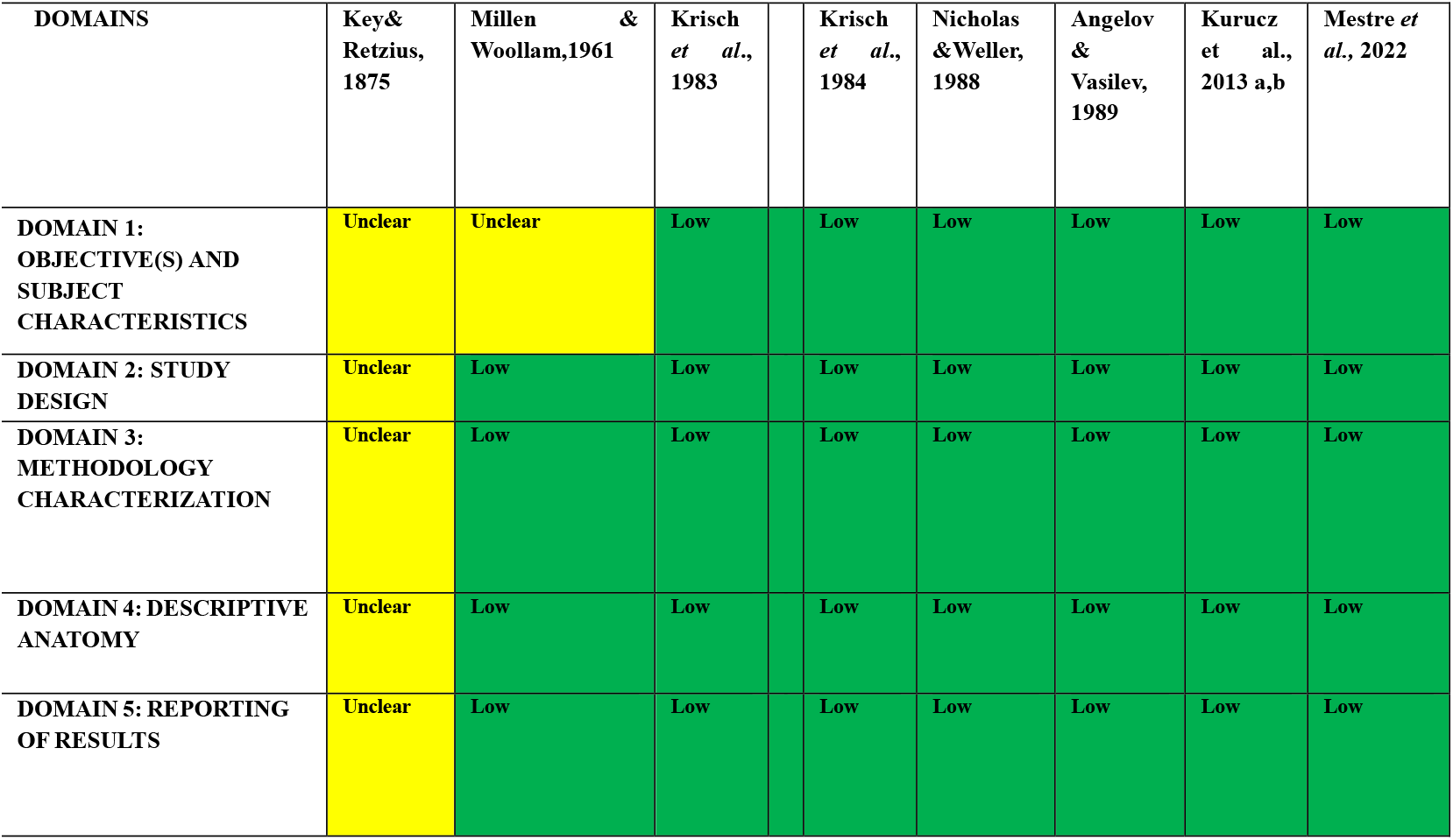
Anatomical Quality Assessment (AQUA) Tool for the quality assessment of anatomical studies included in meta-analyses and systematic reviews.

